# Automated mini-bioreactors reveal the temporal dynamics and multi-omics responses of CRISPRi knockdowns in *Pseudomonas putida*

**DOI:** 10.64898/2026.03.06.709552

**Authors:** Mariana Arango Saavedra, Sara Grassi, Magnus Ganer Jespersen, Catarina Rocha, Viji Kandasamy, Lars Keld Nielsen, Pablo Ivan Nikel, Stefano Donati

## Abstract

Characterizing CRISPR interference (CRISPRi) phenotypes presents a fundamental temporal challenge: pre-existing overabundance of target proteins can mask early silencing, requiring extended growth for dilution, yet prolonged repression rapidly selects for escaper mutants. To resolve this, we integrated a tightly regulated CRISPRi system in *Pseudomonas putida* with an automated mini bioreactor platform operating in turbidostat mode. By maintaining continuous exponential growth, we mapped the exact temporal dynamics of essential gene silencing. We identified a critical observation window between 17 and 27 hours (7–9.5 cell doublings) where repression exerts its maximum physiological impact, directly preceding population takeover by target-site mutated escapers. Applying this workflow to the arginine biosynthesis pathway, multi-omics profiling disentangled transient physiological buffering from long-term mutational events, revealing that *argH* and *argG* knockdowns trigger highly diverse metabolomic perturbations. This scalable framework overcomes batch culture limitations, ensuring precise temporal control for accurate phenotypic characterization and reliable functional genomics.

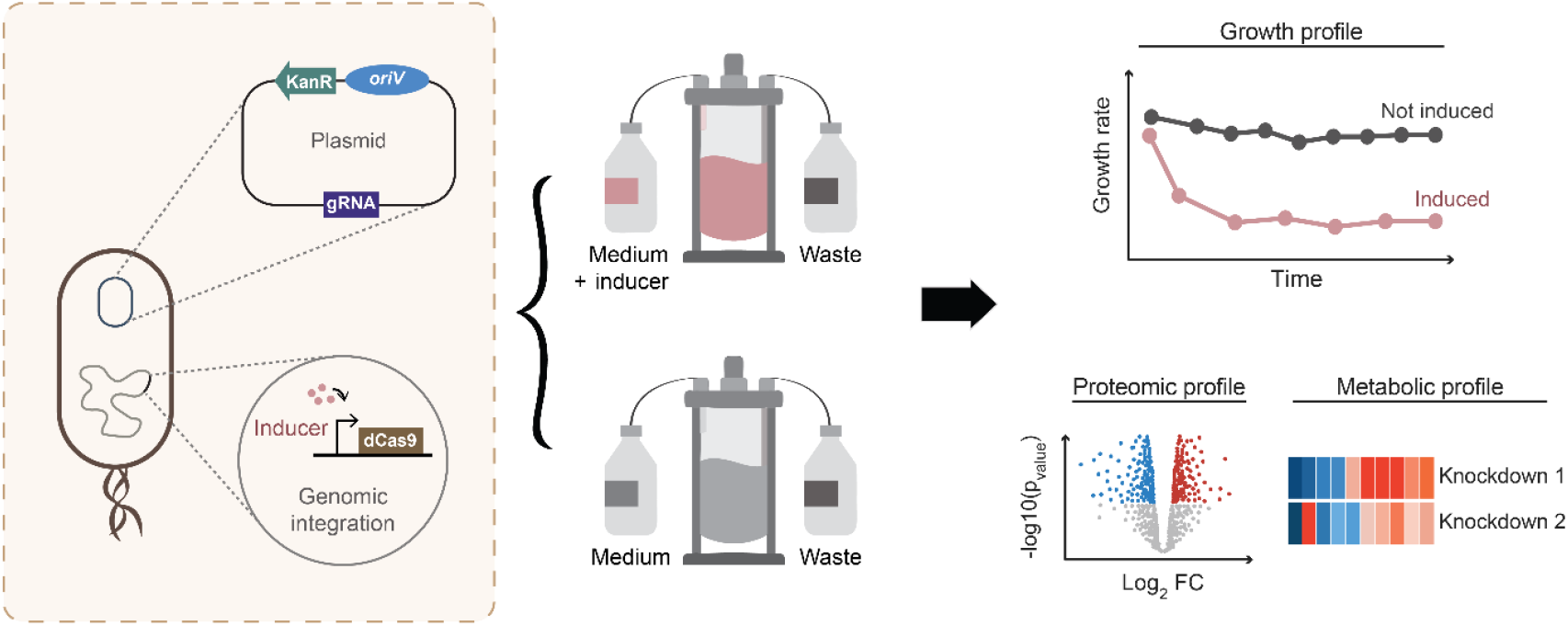

## Introduction

Microbial cell factories are increasingly engineered to produce high-value chemicals and materials from renewable carbon feedstocks, serving as highly efficient platforms for complex, multi-step catalysis^1–3^. However, optimizing these living factories requires a deep understanding of their intricate metabolic networks and regulatory mechanisms. The continuous development of advanced genetic tools has been crucial for dissecting these pathways, guiding metabolic engineering efforts to improve cellular design and enhance targeted physiological traits^4–6^.

A powerful approach for identifying metabolic bottlenecks, uncovering regulatory circuits, and revealing compensatory mechanisms in a cell is through the control of gene expression^7^. CRISPR interference (CRISPRi) offers a simple, easily programmable, inducible, scalable, and non-permanent technique for studying gene repression^8,9^. This method consists of using a catalytically inactive Cas9 (dCas9) that, guided by a single-guide RNA (sgRNA), binds to a specific DNA sequence and prevents RNA polymerase binding or elongation, resulting in transcription inhibition^10^. Although versatile and effective, cells can escape CRISPRi-mediated repression through spontaneous mutations or adaptive responses^11^. This challenge is compounded by the inherent kinetics of CRISPRi. Because the system depends on active transcription during growth, cells entering the stationary phase can obscure repression phenotypes. Furthermore, pre-existing pools of overabundant target proteins can delay or mask observable knockdown effects^12^, requiring researchers to artificially extend exponential growth to dilute existing proteins prior to repression. This creates a fundamental experimental trade-off: extending exponential growth is required to reveal the true knockdown phenotype, but prolonged repression maximizes the selective pressure for escaper mutants to emerge and ultimately overtake the population.

Traditionally, this trade-off is managed by performing manual serial dilutions in batch cultivations to extend exponential growth and dilute existing proteins prior to repression^12,13^. While automated approaches have been implemented, such as the use of multicultivation systems for continuous cultivation of phototrophic organisms in pooled libraries^14,15^ or liquid-handling robots for arrayed libraries and screening workflows^16,17^, these strategies rely on specialized equipment that can be costly, limiting experimental access and throughput. An alternative is represented by recently commercialized affordable 3D-printed mini-bioreactor platforms ^18,19^, which enable users to perform turbidostat or chemostat experiments at a fraction of the cost of other commercial alternatives. By maintaining steady-state physiological conditions, these continuous platforms enable diverse applications, ranging from adaptive laboratory evolution to the precise side-by-side comparison of engineered strains.

In this study, we used mini-bioreactors to investigate the dynamics and stability of CRISPRi in *Pseudomonas putida*, a biotechnological relevant strain characterized by metabolic versatility and high stress tolerance. Using turbidostat cultivations, we identified the optimal number of generations required to reliably detect the effects of repression of essential genes before the emergence of escaper mutants. Additionally, we employed a multi-omics approach to characterize cellular responses to repression of essential genes in the arginine biosynthesis pathway. By combining continuous cultivation with systems-level metabolomic and proteomic resolution, we were able to successfully disentangle transient physiological buffering from permanent mutational escapes. Together, these results establish an inexpensive automated workflow for CRISPRi-based gene repression studies that enables confident interpretation of knockdown phenotypes and is compatible with high-throughput applications.

## Results

### Construction of a tight and inducible CRISPRi system for single-gene knockdowns in *Pseudomonas putida*

To study the effects of single gene repression in *P. putida*, we established a tunable CRISPRi system using a catalytically dead Cas9 (dCas9). The control of gene expression with dCas9 can be compromised by toxicity arising from off-target effects^20^, a problem commonly associated with plasmid-based systems. To mitigate this toxicity, we constructed a *P. putida* MAS strain by integrating the dCas9 cassette into a conserved, non-essential chromosomal region downstream of the *glmS* gene. In this CRISPRi system, expression of dCas9 is controlled by a rhamnose-inducible promoter, whereas the single-guide RNA (sgRNA) is encoded in a plasmid and constitutively expressed (**Fig. 1A**). We determined 3 mM rhamnose to be the optimal inducer concentration that yielded strong gene repression without affecting growth performance (**Supplementary Fig. S1**) and therefore used this concentration in all subsequent induced experiments. Since the efficiency of dCas9 repression largely depends^21^￼, we designed guides to bindr the non-template strand near the start codon (ATG) for all experiments. Additionally, we used non-targeting guides as controls.

**Figure 1.**
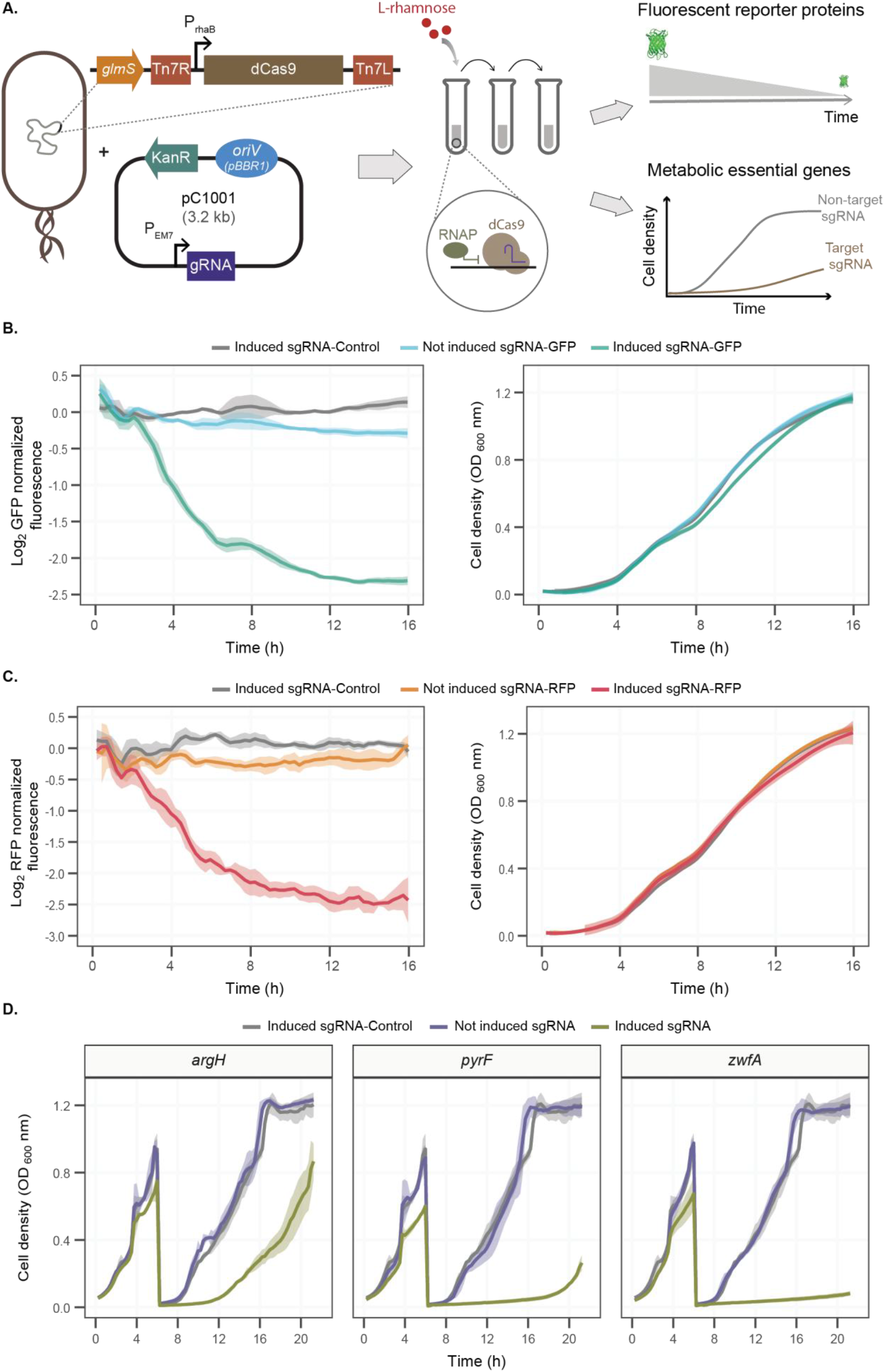
Tunable CRISPRi system in *P. putida*. **(A)** Schematic workflow for establishing a CRISPRi toolset. A rhamnose-inducible dCas9 cassette is chromosomally integrated downstream of the *glmS* conserved region. Fluorescent reporter and essential gene repression are achieved by providing a pC1001 plasmid carrying a targeting sgRNA and inducing dCas9 expression with rhamnose over several dilutions, allowing quantification of decreases in fluorescence or cell density over time. **(B**,**C)** Fluorescence and growth profiles of genomically integrated GFP and RFP knockdowns, respectively. Fluorescence is measured after CRISPRi induction with 3mM rhamnose at time 0 in cells carrying either a fluorescent targeting sgRNA or a non-targeting sgRNA (referred to as sgRNA-Control). Fluorescence is normalized to OD and to fluorescence of an uninduced sgRNA-Control strain. **(D)** Growth profile of strains carrying sgRNA targeting metabolically essential genes or sgRNA-Control under induced and non-induced conditions. CRISPRi was induced with 3 mM rhamnose at time 0. Cultures were re-diluted to fresh medium with or without rhamnose after 6 hours. Data represents means of biological triplicates, with shaded areas indicating standard deviations.

To evaluate the tightness and robustness of the CRISPRi system, we designed plasmids with sgRNAs targeting the expression of fluorescent proteins and essential metabolic genes (**Fig. 1A**). For the repression of fluorescent reporter genes, we generated additional *P. putida* MAS strains by integrating the dCas9 cassette into *P. putida* KT2440 derivatives carrying an additional genome integration of either msfGFP or ^22^. Fluorescence levels of both msfGFP and mRFP decreased exponentially only in the presence of the inducer, while growth remained unaffected (**Fig. 1B** and **1C**). This indicates that the CRISPRi system was robust and that dCas9 expression did not impose a detectable metabolic burden, as uninduced targets and induced controls showed growth profiles similar to those of the induced targeting strains. Fluorescence decreased gradually until stationary phase, consistent with dilution of pre-existing basal protein by growth. The constant fluorescent expression in uninduced cultures with targeted sgRNAs demonstrated minimal CRISPRi system leakage.

To further assess the efficiency and specificity of the system, we chose to target the following metabolic genes: *argH* (argininosuccinate lyase, catalyzing the final step of arginine biosynthesis), *pyrF* (orotidine-5’-phosphate decarboxylase, essential for de novo uracil biosynthesis), and *zwfA* (glucose-6-phosphate dehydrogenase isoform initiating the oxidative branch of the pentose phosphate pathway). We cultivated the strains in chemically defined media (MSM) in which all these genes are essential for growth. We first grew strains for 6 hours in exponential phase under both induced (3 mM rhamnose) and uninduced conditions, followed by cell dilution and continued measurement of optical density (OD_600_ nm) in plate readers for another 14 hours. This allowed us to extend the length of the batch cultures and consequently to extend the length of the CRISPRi experiment (**Fig. 1D**). In induced cultivations, growth defects appeared after ca. 4 hours of induction and became more pronounced following re-dilution. Beyond confirming a strong gene repression and supporting the high specificity and tight regulation of our CRISPRi system, these results underscore the importance of protein dilution by growth for essential genes to reveal the full phenotypic effects of CRISPRi.

### CRISPRi knockdowns in an automated continuous cultivation platform

Due to the growth dependency of CRISPRi experiments, culturing cells in continuous mode can automate and facilitate experimental work. The recent availability of affordable mini-bioreactors can help increase the throughput of such experiments at low cost ^18,19^. Here, we integrated the CRISPRi system into a mini-bioreactor in turbidostat-mode to maintain cells in exponential phase and extend the time of a CRISPRi experiment, further strengthening the knockdown of the target gene (**Fig. 2A**). The setup consisted of mini-bioreactors (Pioreactor), adjusted with automated dosing to perform serial dilutions, maintaining cells in exponential phase throughout the experiments. We set an OD threshold of 0.8 to prevent entry into stationary phase; when cultures reached this threshold, fresh medium was automatically supplied, and excess culture was removed. This experimental approach maintained a constant cell population and enabled monitoring of gene silencing effects over multiple generations.

**Figure 2.**
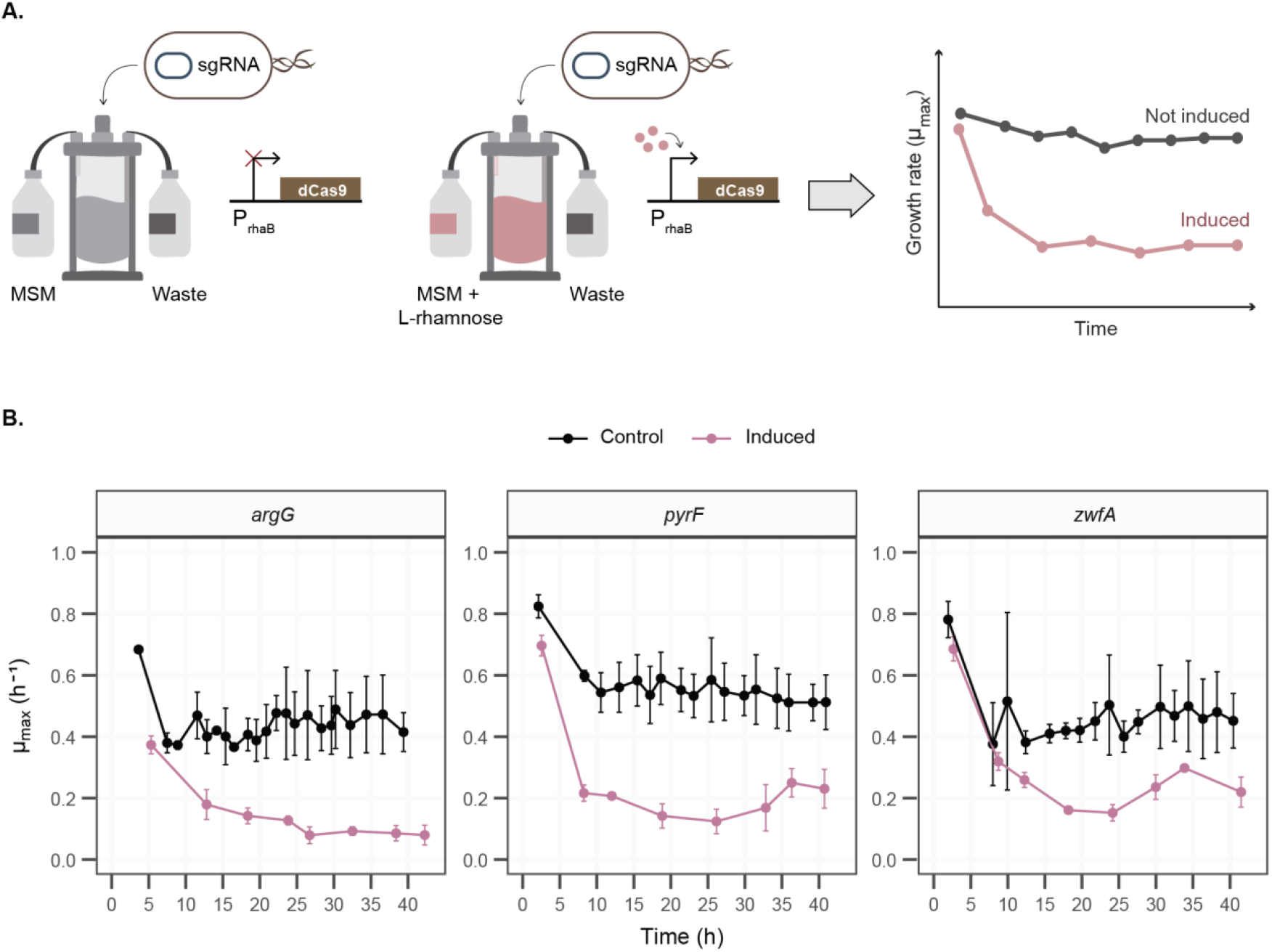
Dynamics of single-gene knockdowns of metabolically essential genes in bioreactors. **(A)** Schematic illustration of the automated bioreactor setup for continuous cultivation of CRISPRi-induced cells. dCas9-integrated cells carrying a targeting sgRNA are either induced with rhamnose or left uninduced and cultivated in a turbidostat. Automated periodic dilutions maintain the cells in mid-exponential growth over extended periods, enabling gradual dilution of basal levels of the targeted protein. Enzyme and metabolite levels can be monitored by sampling at different time points, as indicated by the dashed lines. **(B)** Maximum growth rates of dCas9-expressing cells carrying sgRNAs targeting *argG, pyrF*, and *zwfA*, cultivated in a bioreactor under either uninduced (Control) or rhamnose induced (3 mM) conditions at time 0. Means are shown with error bars indicating the standard deviation of three biological replicates.

We cultivated cells with sgRNAs targeting *pyrF, zwfA, argH*, and *argG* (encoding argininosuccinate synthase, catalyzing the penultimate step in arginine biosynthesis) in 17 mL of chemically defined media (MSM) in which all these genes are essential for growth. For the induced conditions, rhamnose was added from the start of the cultivation. To track the effects of gene repression, we calculated maximum growth rates (µ_max_) for each growth curve between dilution events. Similar to the plate reader experiments, growth rates under induced conditions decreased significantly compared to those under uninduced conditions (**Fig. 2B** and **3A**). Interestingly, the strongest reduction in growth across all sgRNA targets occurred between 17 – 27 hours of cultivation, corresponding to 7 – 9.5 cell-doublings in the induced strains. After this period, growth rates slightly increased.

**Figure 3.**
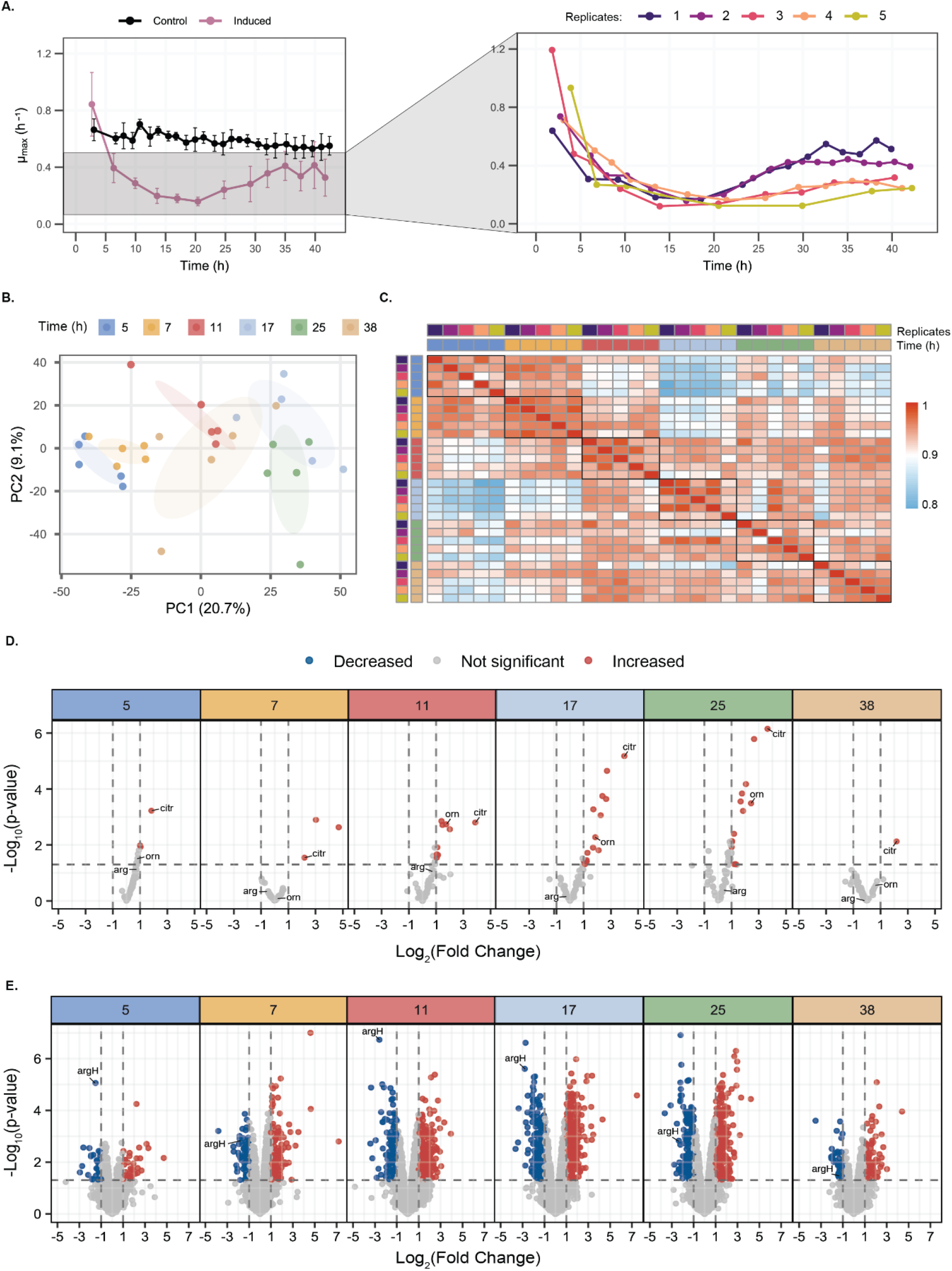
Physiological and molecular responses of the dynamic *argH* CRISPRi knockdown over time. **(A)** Maximum growth rate of *argH* CRISPRi knockdown cultures under induced and uninduced (Control) conditions. The left graph shows mean values from five biological replicates (induced) and three replicates (Control), with error bars representing standard deviations. The right graph shows the distribution of maximum growth rates for individual induced replicates across time points, illustrating differences in growth patterns. **(B)** Principal Component Analysis (PCA) of the proteomic profiles from induced *argH* knockdown samples. Samples cluster according to sampling time, indicating distinct temporal shifts in the proteome during the knockdown response. **(C)** Heatmap of pairwise Pearson correlation coefficients between biological replicates of the induced *argH* knockdown across time points. Colors indicating time and replicate identity follow those shown in graphs (A) and (B). **(D, E)** Volcano plots showing changes in metabolite (D) and protein (E) abundance in the induced *argH* knockdown compared to the uninduced control across cultivation time points (5 – 38 h). Colors indicating time follows those shown in (B). Each time point highlights significantly changed metabolites (|log_2_(fold change)| > 1 and p-value < 0.05) or proteins (|log_2_(fold change)| 1 and adjusted p-value < 0.05 using the Benjamini-Hochberg method), showcasing the progression of metabolic and proteomic shifts over time.

### Cellular responses to the temporal dynamics of CRISPRi in continuous cultivations

Given the previously observed growth recovery patterns, we asked whether the underlying physiological perturbations persisted and how they might evolve over a prolonged knockdown period. To this end, we examined the cellular responses of *argH* knockdown strains over 42 hours of induction, with 5 replicate mini-bioreactor cultivations for the induced condition and 3 replicates for the control un-induced condition (**Fig. 3**). As previously described, the knockdown effect appeared to weaken after 20 hours of culture, and inspection of individual replicates revealed distinct recovery trajectories (**Fig. 3A**). To determine whether these recoveries resulted from spontaneous mutations in the CRISPRi system, we performed whole-genome and plasmid sequencing of population samples at the end of the experiment. Sequencing results showed no mutations in the plasmid vector carrying the sgRNA or in the integrated dCas9 cassette. However, three specific point mutations were identified in the *argH* target region of the chromosome in three of the five replicates, each present at low frequency within the population (**Supplementary Table S4** and **Supplementary Fig. S2**). The identified mutations were located near the start codon, with one being synonymous and two non-synonymous. Replicates carrying non-synonymous mutations (replicates 1 and 2) exhibited the largest increase in growth rate, whereas slow-recovering replicates had either a mixture of synonymous and non-synonymous mutations (replicate 4) or no mutations in the *argH* region (replicates 3 and 5). This indicated that escaper mutants able to circumvent the CRISPRi system could overtake the culture in the mini-bioreactors, albeit after several hours of continuous growth.

We then verified whether these mutations could be affecting the CRISPRi system and consequent gene expression responses. We monitored the metabolome and proteome of the replicates at multiple time points during induction. Principal component analysis (PCA) of the proteome, showed distinct clusters and time-dependent patterns (**Fig. 3B**). Notably, samples from the latest time point (38 h) were more dispersed, revealing greater variability among replicates and corresponding with divergent growth behaviors of the different replicates. Over time, samples generally separated further in the PCA space, with those from 17 and 27 h being the most distant, and those from 38 h appearing closer to the earliest time points (5 and 7 h). High correlations among samples from the same time points reinforced the experiment’s reproducibility and revealed gradual divergence over time (**Fig. 3C**). Of particular interest are samples at 17 h, which showed the lowest correlations with earlier time points, suggesting that the strongest CRISPRi effect on the proteome occurred around this time, after which cells began restoring their initial proteome activity. Correlations of the uninduced control replicates exhibited minimal change over time, denoting that the observed changes in the induced conditions were not intrinsic to the turbidostat cultivation setup (**Supplementary Fig. S3**). As time progressed, both the metabolome (**Fig. 3D**) and proteome (**Fig. 3E**) of induced cells were progressively more perturbed, before eventually reverting to a less perturbed state at 38 h. A similar trend was observed when examining the changes in individual proteins and metabolites within the arginine pathway, including the target of interference ArgH, at its lowest abundance at the 17h timepoint (**Supplementary Fig. S4** and **S5**). While the target of interference ArgH was strongly decreased, levels of dCas9, of the transcriptional activator RhaS, of the transcriptional activator RhaR and the kanamycin-resistant protein KanR remained relatively stable over time (**Supplementary Fig. S6**). The consistently low abundance of dCas9 in the control samples further confirmed the tight control of the expression system. NADH/NAD^+^ and NADPH/NADP^+^ ratios remained stable and the adenylate energy charge (AEC) remained above 0.7 throughout the experiment (**Supplementary Fig. S7**), indicating that cells-maintained ATP balance and overall energy homeostasis during the experiment.

Taken together, the observed dynamics confirmed that the strongest effect of CRISPRi occurred between 17 and 27 h for the *argH* knockdown, before the escaper mutants could outgrow cells still experiencing the CRISPRi knockdown.

### Knockdowns of genes in the same biosynthetic pathway cause a different response at the metabolome level

Next, we analyzed whether perturbing closely related genes of the same pathway can lead to different responses. We focused on the arginine pathway, perturbing expression of *argH* and *argG*, which encode for sequential enzymes in the pathway (**Fig. 4**). Arginine is synthesized via the acetylated ornithine pathway, whose final step is catalyzed by ArgH, and is degraded through catabolic pathways that convert it back to ornithine, glutamate, and other intermediates, allowing the cell to recycle nitrogen and carbon^23^. An important regulator of this network is the transcriptional factor ArgR, which acts as a negative regulator by repressing the expression of the pathway in response to high intracellular arginine levels^25,26^.

**Figure 4.**
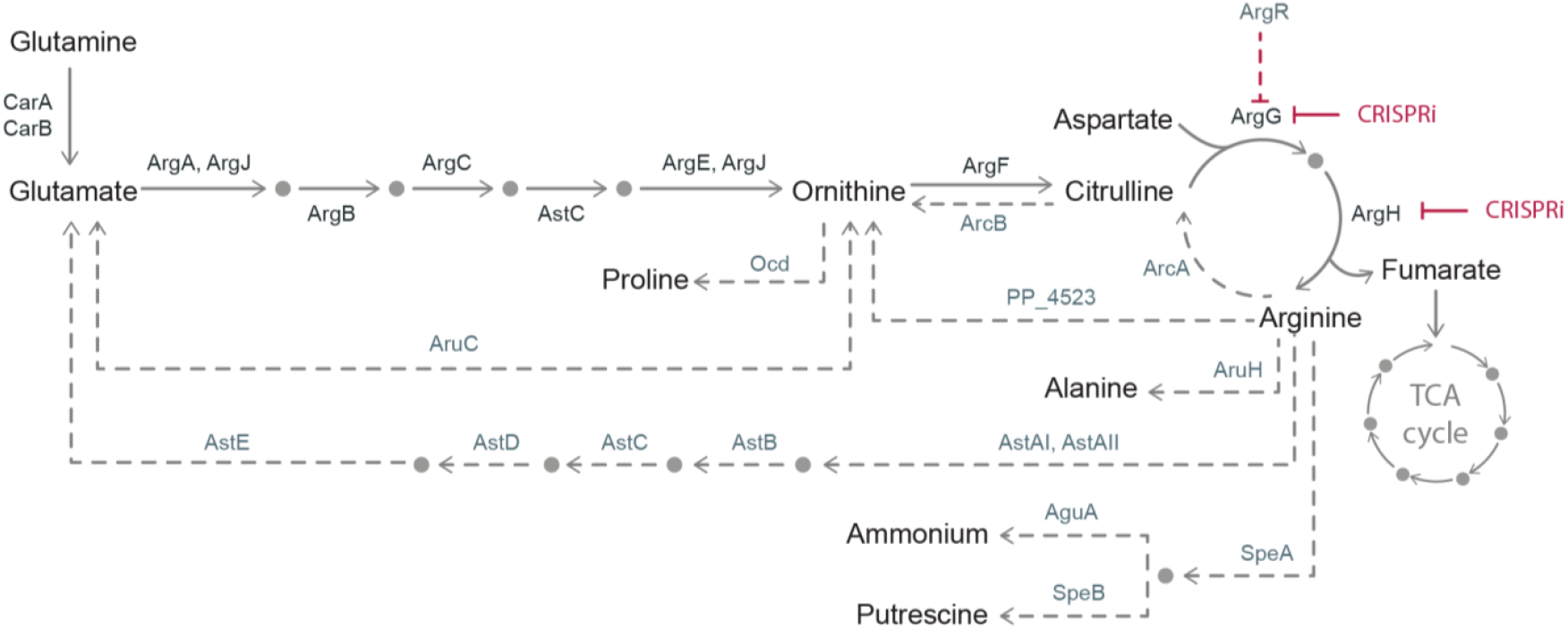
Arginine biosynthesis and degradation pathways. Model illustrating proteins and selected metabolites involved in arginine metabolism in *P. putida*, with biosynthesis shown by solid lines and degradation by dashed lines. Glutamate, derived from glutamine, serves as the starting point for arginine de novo synthesis via the acetylated ornithine pathway, where glutamate is sequentially converted to ornithine, then to citrulline and finally to arginine. Arginine can be catabolized through multiple routes. The ADI pathway, involving ArcA and ArcB, converts arginine to ornithine via citrulline. Alternative routes include the conversion back to glutamate through the AST catabolic pathway (involving AstAI, AstAII, AstB, AstC, AstD, and AstE), conversion to alanine, or to putrescine and ammonium. Ornithine produced from arginine degradation can further feed into proline or glutamate biosynthesis. Arginine biosynthesis is negatively regulated by ArgR via feedback inhibition on ArgG, indicated by the red dashed line. The targets of the CRISPRi are shown by solid red lines.

Considering the previous observations of the temporal dynamics of our system, we analyzed metabolomic samples after 17 hours of continuous CRISPRi induction (**Fig. 5A**). In particular, the *argH* knockdown caused a broader and stronger metabolic disturbance, characterized mainly by increased metabolite levels. Nonetheless, accumulation of the arginine biosynthesis intermediates aspartate (asp), ornithine (orn), and citrulline (citr) was observed in both knockdowns. Similarities between these two knockdowns were also encountered for metabolites involved in the pentose phosphate pathway, glycolysis, gluconeogenesis, and TCA cycle. However, a noticeable difference between the two knockdowns was that *argH* showed increased levels of metabolites involved in nucleotide metabolism (adenine (ade), adenosine (adn), camp, cmp, ctp, datp, cdcp, cdmp, dgdp, dtmp, ump, uracil (ura), uridine (uri), and utp) and amino acid metabolism (alanine (ala), glycine (gly), methionine (met), phenylalanine (phe), serine (ser), threonine (thr), tryptophan (trp), and tyrosine (tye)).

**Figure 5.**
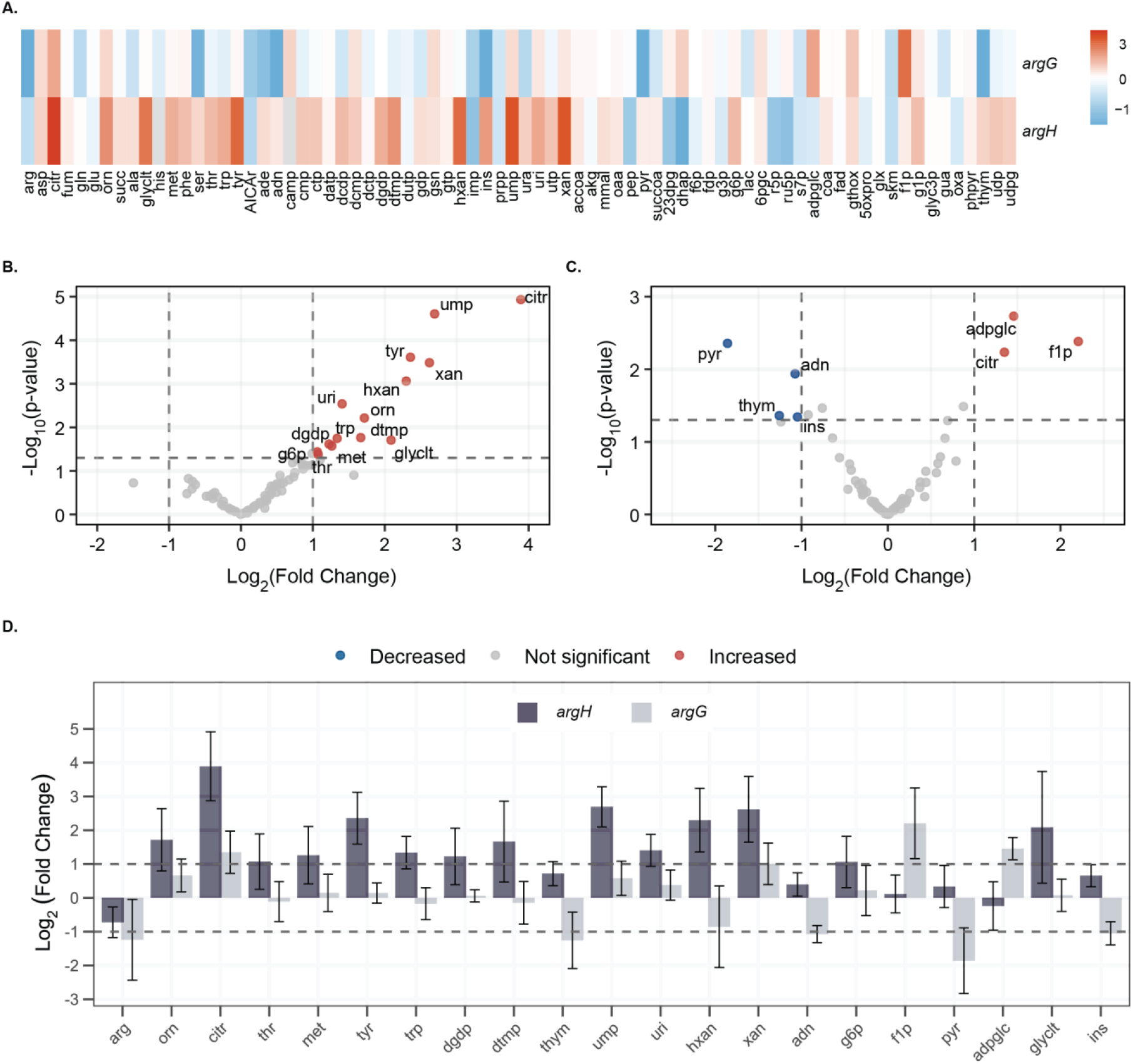
Intracellular metabolomic responses of *argH* and *argG* CRISPRi knockdowns. **(A)** Heatmap with color gradient showing mean of log_2_(fold change) in the relative abundance of 76 intracellular metabolites for *argH* and *argG* knockdowns compared to their respective uninduced controls. **(B, C)** Volcano plots showing log_2_(fold change) versus -log_10_(p-value) for *argH* (B) and *argG* (C) knockdowns relative to their corresponding uninduced controls. Each dot represents an individual metabolite. Dashed vertical lines indicate the threshold for |log_2_(fold change)| > 1, and the horizontal dashed line marks the significant threshold (p-value < 0.05). **(D)** Comparison of significantly changed metabolites identified in the volcano plots for *argH* and *argG* knockdowns. Dashed horizontal lines indicate the significance threshold (log_2_(fold change) 1 or < -1). Data represent mean values from five biological replicates for *argH* and three biological replicates for *argG*, with error bars indicating standard deviations.

To better understand the distinct cellular responses between the two knockdowns, we examined metabolites that showed statistically significant changes in relative abundance. Only a few metabolites were significantly affected in the *argG* knockdown (**Fig. 5C**), whereas *argH* showed a greater number of significantly altered metabolites (**Fig. 5B**), all of which exhibited increased abundances, indicating a more global metabolic imbalance. The accumulation of ornithine and citrulline was more pronounced in *argH* knockdown, while arginine, the final product of the pathway, had a minor decrease in abundance for both knockdowns, with no significant changes (p-value > 0.05)(**Fig. 5D**). This moderate decrease in arginine suggests maintenance of the intracellular pool via residual enzymatic activity and/or high turnover of this metabolite, or through compensatory metabolic routes. The concentration of thymine (thym), hypoxanthine (hxan), pyruvate (pyr), and inosine (ins) significantly decreased, but only in the argG strain. The observed differences indicate that while metabolites closely related to the different metabolic bottlenecks were perturbed similarly, the perturbation propagated differently, leading to nucleoside accumulation in the *argH* knockdown and depletion in *argG*.

### Proteomic characterization of the *argH* CRISPRi knockdown

When analyzing the global proteomic profile after 17 hours of CRISPRi induction to assess the broader cellular response to *argH* knockdown (**Fig. 6A**), we observed both expected and compensatory changes. Among these, there was a ca. 7-fold reduction in ArgH, further confirming the effectiveness of the CRISPRi system. Upregulation of enzymes involved in the previous biosynthesis steps of the pathway was also observed. In contrast, other enzymes, such as CarA and CarB (the small and large subunits of the carbamoyl-phosphate synthase that supply carbamoyl phosphate to the early steps of the pathway) were downregulated. Collectively, these changes reflect the mechanisms that the cell activates in response to depletion of ArgH and arginine, possibly aiming at restoring a flux through the pathway.

**Figure 6.**
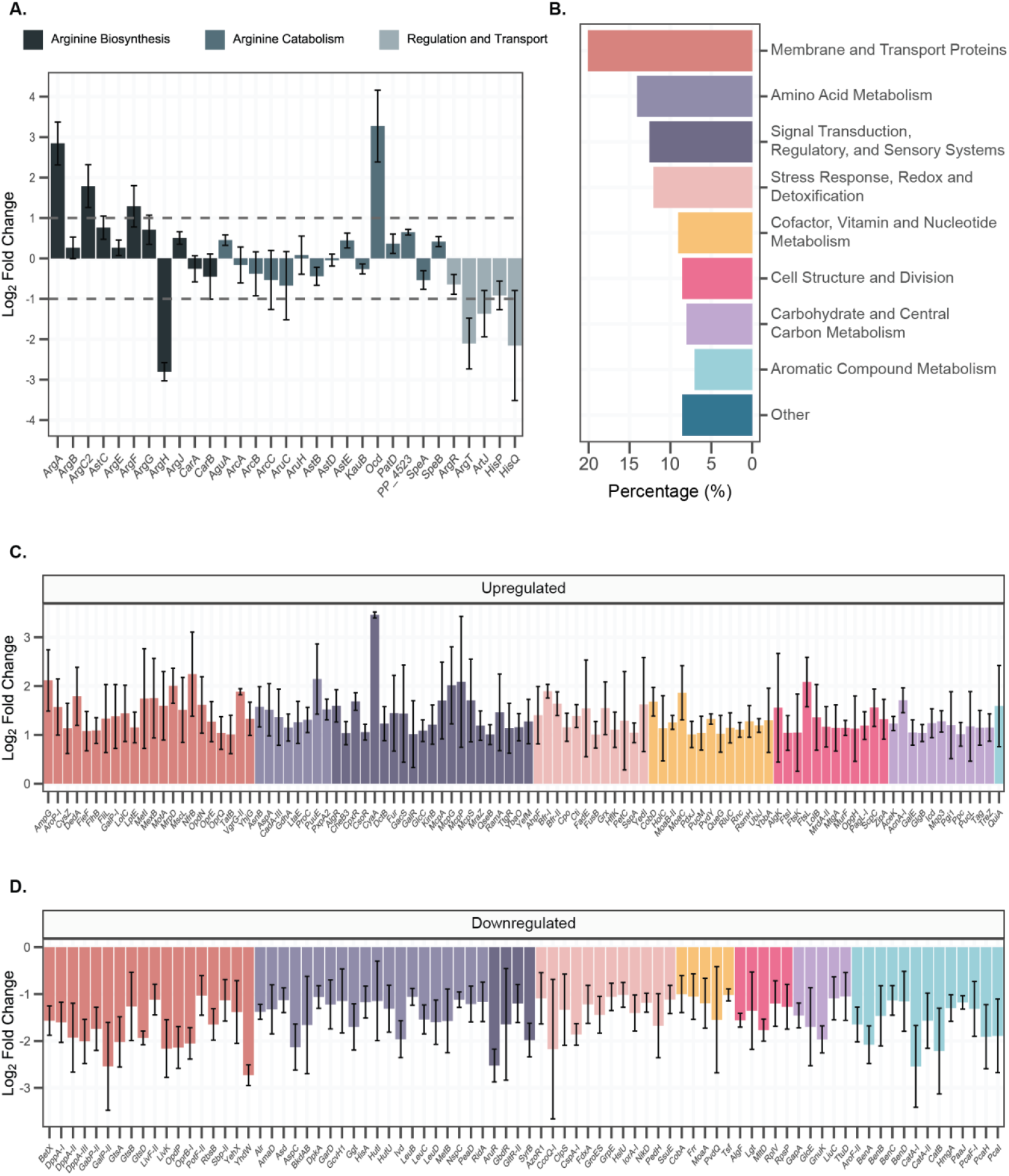
Proteomic profile of the *argH* CRISPRi knockdown. **(A)** Changes in abundance of proteins directly involved in arginine metabolism after 17 hours of *argh* knockdown, shown as log_2_(fold change). Proteins are grouped according to their function (biosynthesis, catabolism, and transport and regulation). Dashed horizontal lines indicate significance threshold (log_2_(fold change) > 1 or < -1). **(B)** Proteins showing significant differential expression (p-values of < 0.05 and |log_2_(fold change)| >1) and with full functional annotation (i.e., with assigned gene names) were classified into functional categories based on their primary biological processes. The percentage indicates the proportion of these significantly changed proteins (n = 199 out of 3664 total quantified proteins) that fall within each functional category. **(C**,**D)** Subset of significantly upregulated and downregulated proteins (p-values of < 0.05 and |log_2_(fold change)| >1), respectively. The color code follows that shown in panel (B). Proteins within the ‘Other’ functional category are displayed in Supplementary Fig. S8. Data on bar plots represent mean values from five biological replicates, and error bars indicate standard deviation.

Enzymes involved in arginine catabolism were generally downregulated, with the exception of Ocd (ornithine cyclodeaminase) and the enzyme encoded by PP_4523, which both showed increased levels. Together with the prominent accumulation of ornithine, this suggests an activation of a feedback route that channels arginine back to ornithine, possibly as part of a nitrogen recycling mechanism. On the other hand, the amino acid transport proteins ArgT, ArtJ, HisP, and HisQ were downregulated, likely as a cellular strategy to limit extracellular uptake in response to intracellular imbalance of their corresponding substrates. Regarding the transcriptional regulator ArgR, its abundance decreased moderately, while its target gene *argG* was upregulated, confirming what has been previously reported on the strong negative regulatory effect that ArgR exerts on *argG*^25,27^.

The proteomic response was localized, with only ca. 5.4% of all detected proteins showing significant changes in abundance (|log_2_(fold change)| >1) (**Supplementary Table S5**). We then grouped these proteins into functional categories based on their biological roles (**Fig. 6B**) and observed that the most impacted categories were membrane and transport proteins, amino acid metabolism, signal transduction, and regulatory and sensory systems, which supports our previous observations. Other affected proteins included those involved in ribosome recycling, elongation, and other general cellular activities, reflecting the slow growth state of the cell. A small subset encompassed hypothetical proteins or proteins with poorly characterized functions (**Supplementary Fig. S8**).

The distribution of categorized proteins between upregulation (**Fig. 6C**) and downregulation (**Fig. 6D**) was roughly equal for those involved in stress response, redox and detoxification, and membrane and transport proteins. Proteins in the signal transduction, regulatory and sensory systems, as well as in the cell structure and division, and carbohydrate and central carbon metabolism categories were mostly upregulated, indicating a global physiological response to cell stress. Conversely, most proteins involved in amino acid metabolism and aromatic compound metabolism were downregulated. Downregulation of proteins involved in amino acid metabolism may reflect metabolic reprogramming in response to perturbations in arginine biosynthesis (**Fig. 5D**), whereas the downregulation of proteins involved in aromatic compound metabolism likely reflects carbon catabolite repression (CCR), which is activated when cells are grown on glucose as sole carbon source to prioritize its consumption while repressing genes involved in lesser preferred carbon sourced like aromatic compounds^28,29^.

## Discussion

In this study, we expanded the CRISPRi toolset in *P. putida*^13,30–34^ by constructing and characterizing a tight and robust CRISPRi system consisting of a dCas9 genomically integrated and controlled by a rhamnose-inducible promoter (*rhaBAD*). This promoter, native to *Escherichia coli*, has been widely used in plasmid-based systems across a broad range of bacteria due to its tight expression control and its high tunability properties, including its linear response to the inducer^35–40^. Additionally, rhamnose is non-toxic and non-metabolizable in *P. putida*, allowing it to remain stable in the media and precisely control the level of gene activation^40^ by enabling sustained induction. This was demonstrated with the knockdown of both fluorescent reporters and essential genes, where minimal leakage was observed. Beyond its applicability to batch cultures, we demonstrated that implementing CRISPRi in continuous cultivation using automated and affordable mini-bioreactors. As a result, we were able to deplete target proteins from a basal level to a critical level through dilutions and observe the most prominent gene silencing effect after 17-27 hours of cultivation. This corresponds to ca. 7-9.5 cell-doublings of the induced strain and indicates that substantial dilution of pre-existing protein is required before physiological effects become apparent. In a pooled CRISPRi screen of *E. coli* metabolism, it was shown that growth phenotypes emerged with a response time ranging from 4 up to 16 hours after induction^12^. This suggests that performing CRISPRi screens for around 7-9.5 doublings should suffice to identify growth-limiting genes as well as to enforce strong cellular responses also in *P. putida*, before the onset of escaper mutants that might mask these effects.

Another aspect of using continuous cultivations includes maintaining the homogeneity of cell populations in a metabolically active state^41^, which facilitates the acquisition of samples for omics studies focused on characterizing cell physiology at a molecular level and ensuring experimental reproducibility^42^. Cells exposed to stressful conditions can rapidly adapt and alter their molecular responses to overcome the perturbation, as shown in the present study by the dynamic proteomic and metabolomic shifts observed following *argH* knockdown. This is particularly important in the context of CRISPRi for systematic phenotypic analysis of essential genes, where examining the molecular effects of gene knockdowns at an inappropriate time (either too early, before silencing is fully established, or too late, after adaptive responses have begun or escaper mutants have appeared), can lead to underestimation or misinterpretation of results.

By using our CRISPRi system to silence *argG* and *argH*, we were able to examine the metabolic and proteomic activity of the cells in response to perturbations in the arginine pathway. This confirmed the essentiality of both genes, previously demonstrated by the inability of *argH* and *argG* full knockouts to grow on minimal media in the absence of arginine^43^. In our study, metabolomic profiling of *argH* and *argG* knockdowns enabled a comparative assessment of pathway-level perturbations caused by targeting distinct steps in arginine biosynthesis. This showcased that closely related perturbations at the pathway level can propagate in very different directions at the metabolome level, leading to nucleoside accumulation in the *argH* knockdown strain and depletion in the *argG* knockdown strain.

In conclusion, the automated CRISPRi system established in this study is a suitable tool for the systematic study of metabolic pathways using single-gene knockdowns combined with an omics approach. As a proof of concept, we characterized metabolome and proteome reallocation over time of induced knockdowns of relevant proteins in the arginine biosynthesis pathway. This workflow was also effective for the detection of compensatory pathways and metabolic bottlenecks, confirming that CRISPRi escapes are primarily caused by physiological adaptations rather than mutational events. Furthermore, we provide a platform that can be implemented in high-throughput and dynamic studies, including large genome-wide CRISPRi screens for functional genomics. Notably, future studies should consider the transient responses of continuous knockdowns and the physiological buffering capacity of cells when assessing the effects of gene repression.

## Materials and Methods

### Strains and culture media

*E. coli* chemically competent NEB 10-beta (New England BioLabs Inc.) cells were used as hosts for standard cloning, while *E. coli* electrocompetent OneShot Top10 (Thermo Fisher Scientific Inc.) cells were used when high transformation efficiency was required. For conjugation by four-parental mating, *E. coli* Dh5α cells served as donor and helper strains. *Pseudomonas putida* KT2440 and its msfGFP and mRFP-integrated derivatives were used as parent strains for engineering purposes. All strains and plasmids used are listed in Supplementary Table S1 and Table S2, respectively.

Lysogeny broth (LB) medium (10 g/L tryptone, 5 g/L yeast extract, and 5 g/L NaCl) was used for routine cultivation of all bacterial strains with shaking at 250 rpm. *E. coli* strains were grown at 37°C, while *P*. putida strains were grown at 30°C. Vogel-Bonner minimal agar medium (VBMM) (3 g/L trisodium citrate, 2 g/L citric acid, 10 g/L K_2_HPO_4_, 3.5 g/L NaNH_4_PO_4_·4H_2_O, 15 g/L agar, 0.24 g/L MgSO_4_·7H_2_O, and 14.6 mg/L CaCl_2_·2H_2_O; pH 7) was used to counterselect *E. coli* after conjugation.

Knockdown growth experiments were conducted in minimal salt medium (MSM) medium (3.88 g/L K_2_HPO_4_, 1.63 g/L NaH_2_PO_4_, 2 g/L (NH_2_)_2_SO_4_, and 0.1 g/L MgCl_2_·6H_2_O; pH 7) supplemented with trace elements (10 mg/L EDTA, 2 mg/L ZnSO_4_·7H_2_O, 1mg/L CaCl_2_·2H_2_O, 5 mg/L FeSO_4_·7H_2_O, 0.2 mg/L Na_2_MoO_4_·2H_2_O, 0.2 mg/L CuSO_4_·5H_2_O, 0.4 mg/L CoCl_2_·6H_2_O, and MnCl_2_·2H_2_O) and 20 mM glucose. The medium for knockdown experiments was additionally supplemented with 50 µg/mL kanamycin for all conditions, and with 3 mM rhamnose only for the induced conditions. For plasmid and strain construction, antibiotics were added at the following concentrations when required: 25 µg/mL chloramphenicol, 100 µg/ mL ampicillin, 50 µg/mL kanamycin and 20 µg/mL gentamycin.

### Strain construction

*P. putida* MAS strains were constructed by genome integration of the dCas9 expression cassette using the mini-Tn7 transposon system^44^. The cassette consisted of a catalytically dead Cas9 from *Streptococcus pyogenes* (mutations D10A and H840) under the control of the L-rhamnose inducible promoter system (P_rhaBAD_). The pJM220_R1 suicide vector ^45^ harboring by inserting the *SpdCas9* gene from pMCRi downstream of the rhamnose promoter in pJM220. pJM220_R1 was delivered to *P. putida* KT2440 and derivative fluorescent strains through four-parental conjugation mediated by two helper *E. coli* strains carrying the plasmids pRK600 and pUX-BF13. Overnight cultures of the donor, helpers, and *P. putida* recipient strains were grown in LB and combined in a 1:1 OD_600_ ratio. The mixture was centrifuged at 7,000 x g for 2 minutes, washed twice with 1 mL of pre-warmed LB, and resuspended in 30 µL of LB. The cell suspension was spotted onto a pre-warmed LB agar plate containing a 0.45 µm cellulose nitrate membrane filter (Whatman PLC). The plate was incubated at 37°C for 16 hours, after which the filter was transferred to an Eppendorf tube and washed with sterile 0.9 % NaCl to dislodge conjugant cells. 100 µL of the resuspended cells were plated onto VBMM agar plates containing gentamycin and incubated at 30°C overnight. An additional passage of grown colonies was performed on selective VBMM agar plates to remove residual *E. coli* cells.

### Construction of sgRNA-carrying vector

The plasmid pC1001 was constructed using uracil USER cloning^45^ by replacing the *chnR-PchnB* cargo of pSEVA2311 with the sgRNA cargo from pMCRi, which consists of the EM7 promoter, a non-targeting sgRNA region, and sgRNA scaffold. Cloning fragments were amplified using Phusion U Hot Start Polymerase (Thermo Fisher Scientific Inc.) and subsequently treated with *DpnI* restriction enzyme (Thermo Fisher Scientific Inc.) for 8 hours. The treated PCR products were purified using the NucleoSpin Gel and PCR Clean-up Kit (Macherey-Nagel GmbH & Co.), and 100 ng of each fragment was mixed with 1 µL of USER enzyme (New England BioLabs Inc.) in a final volume of 12 µL. The mixture was incubated at 37°C for 25 min, then at 25°C for 25 min, and finally at 10°C for 15 min. Afterwards, 2 µL of the resulting cloning reaction was transformed into 25 µL of chemically competent *E. coli* cells and plated onto LB agar medium supplemented with 50 µg/mL kanamycin.

Replacement of the sgRNA targeting region was performed by digesting the pC1001 plasmid with the Eco31I (BsaI) restriction enzyme (Thermo Fisher Scientific Inc.), followed by dephosphorylation with FastAP Alkaline Phosphatase (Thermo Fisher Scientific Inc.) and ligation of the respective targeting spacers. Forward and reverse strands of the targeting spacers, which were designed with homology arms to the linearized plasmid, were ordered as single-stranded oligonucleotides from Integrated DNA Technologies Inc. Annealing and phosphorylation were carried out using 1 µL of oligonucleotides (100 µM), 1 µL of T4 Polynucleotide Kinase (Thermo Fisher Scientific Inc.), and 1 µL of T4 Ligase Buffer (Thermo Fisher Scientific Inc.). The mixture was incubated as follows: 37°C for 30 min, 95°C for 4 min, and 70 cycles of 12-second temperature decrements, decreasing by 1°C per cycle. Ligation was performed using 1 µL of T4 DNA ligase (Thermo Fisher Scientific Inc.) and a 1:28 molar ratio of spacer to plasmid. The mixture was incubated at 22°C for 30 min, after which 4 µL were transformed into 50 µL of chemically competent *E. coli* cells and plated onto selective LB agar medium. Positive colonies were verified by colony PCR using OneTaq Quick-Load Master Mix (New England BioLabs Inc.). The resulting plasmids were extracted with the NucleoSpin Plasmid EasyPure kit (Macherey-Nagel GmbH & Co.) and transformed into *P. putida* MAS electrocompetent cells, which were prepared by pelleting overnight cultures in LB at 13,000 rpm for 2 minutes and resuspending them twice with 2 mL 300 mM sucrose. 50 ng of the isolated plasmids were mixed with 50 µL of freshly prepared electrocompetent cells into 2 mm gap cuvettes and electroporation was performed with a Gene Pulser Xcell Electroporation System (Bio-Rad Laboratories Inc.) set at 2.5 kV, 25 µF capacitance, and 200 Ω resistance. Oligos used in this study are listed in Supplementary Table 3.

### Cultivation conditions for single CRISPRi knockdowns

Validation of the CRISPRi system was initially performed by measuring the optical density (OD_600_) of the *argH, argG, pyrF, zwfA* knockdown strains, as well as OD_600_ and fluorescence for the m*sfGFP* and *mRFP* knockdown strains using a BioTek Synergy H1 microplate reader (Agilent Technologies Inc.). Single colonies were first inoculated into 5 mL of selective LB medium in culture tubes and incubated overnight. The overnight cultures were then diluted 1:100 into 5 mL of selective MSM medium in culture tubes and incubated overnight. These pre-cultures were diluted to an initial OD_600_ of 0.05 in 2 mL of selective MSM medium, with and without rhamnose. After six hours of incubation, the cultures were further diluted to an OD_600_ of 0.05 in 200 µL of selective MSM medium (with and without rhamnose) in 96-well microtiter plates. To prevent evaporation, 30 µL of mineral oil was added to each well. The plates were incubated in the microplate reader with continuous orbital shaking, and OD_600_ and/or fluorescence was measured every 15 minutes. msfGFP and mRFP fluorescence were measured using excitation/emission wavelengths of 485/528 and 588/633, respectively. Each knockdown strain was cultured in biological triplicates.

Dynamics of the *argG, argH, pyrF* and *zwfA* knockdown strains were evaluated in continuous cultivations using 20 mL turbidostats filled to a working volume of 17 mL (Pioreactor Inc.). Single colonies were inoculated in 2 mL of selective LB medium and incubated overnight. A 1:100 dilution of the overnight cultures was used to start pre-cultures in 5 mL of selective MSM medium in culture tubes, which were also incubated overnight. These pre-cultures were diluted to an initial OD_600_ of 0.05 in selective MSM medium, with and without rhamnose. The target OD_600_ for the turbidostat was set to 0.8; once reached, 9 mL of culture volume was exchanged. The cultures were stirred at 1100 rpm and run for 42 hours. For the *argH* and *argG* knockdown, 1 mL of culture at OD600_nm_ 0.8-0.9 were collected for metabolomics, while 2 mL of culture was sampled for proteomics in the *argH* knockdown. Samples were taken after 5, 7, 11, 17, 25, and 38 hours of cultivation. Cell-doublings for the induced knockdowns during continuous cultivation were calculated using the following equation:

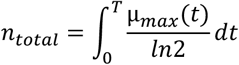

where *n*_total_ corresponds to the total number of cell generations, µ_max_(*t*) is the maximum growth rate (h^-1^) for the dilution interval at time *t*, and T is the total cultivation time (hours).

Samples for whole genome sequencing and plasmid extraction were collected at the end of the experiment.

### Intracellular Metabolite Quantification

Collected samples were prepared by fast filtration through a polyvinylidene fluoride (PVDF) membrane (0.45 µm) under vacuum. Filtered cells were immediately lysed in cold 40:40:20 acetonitrile:methanol:water containing 0.1% formic acid buffer. Dried samples were reconstituted in 90 µL of LC-MS grade water, and isotope diluted samples were prepared by adding 10 µL of a ^13^C labeled internal standard. All cell extracts were analyzed by LC-MS/MS using a SCIEX Exion LC coupled to a SCIEX tripleQuad 6500+ (AB SCIEX), as described in Rocha et al.^46^ Extracted ion scans were processed in SCIEX OS 3.4.5, and the ^12^C:^13^C abundance ratio was calculated for each metabolite. Metabolite abundances were normalized to the OD_600_ at the time of sampling. Quality controls and carryover checks were performed within each batch.

### Protein Quantification

Cells were harvested by centrifugation at 10,000 x g for 10 min at 4°C and the pellets were stored at -20°C until further processing. Frozen pellets were thawed on ice, and the following steps, which included cell lysis, protein quantification, enzymatic digestion, and peptide clean up, were performed following a protocol optimized for HPLC-MS based cytosolic proteomics in gram negative bacteria^47^. Protein concentrations were quantified using a bicinchoninic acid (BCA) assay, and 1 µg of digested peptides were analyzed.

Total proteome was analyzed using data-independent acquisition (DIA) on a Q-Exactive Orbitrap mass spectrometer coupled to Proxeon nanoLC EASY1000 system. The proteome composition was quantified using the standard-free total protein approach (TPA), which calculated the relative abundance of each protein based on its ratio to the sum of all identified proteins^48^. Raw DIA MS files were processed in a library-free workflow using the autoprot pipeline^48^ (available on GitHub: https://github.com/biosustain/autoprot), with peptide and protein identification performed against the reference for *Pseudomonas putida* proteome (Proteome ID UP000000556). In addition, proteins of interest not included in the reference proteome (dCas9, KanR, RhaS and RhaR) were manually added to ensure accurate mapping and quantification.

### Plasmid and Whole Genome Sequencing

Genomic DNA was extracted using the GeneJET Genomic DNA Purification Kit (Thermo Fisher Scientific Inc.). Quality and concentration of the purified DNA were estimated using a Qubit 2.0 Fluorometer and NanoDrop 200/200c Spectrophotometer (Thermo Fisher Scientific Inc.). Samples were sent to Novogene Co. Ltd for library preparation and paired-end (150 bp) sequencing on an Illumina NovaSeq X Plus platform. Initial data quality control of raw reads was done with FASTQC. Reads were then aligned and analyzed for single-nucleotide polymorphisms (SNPs), insertions, deletions, and structural variants using Breseq v0.19.0, with the reference genome assembly for *P. putida* KT2440 (NCBI ID: GCF_000007565.2) used as the mapping template.

Plasmids were isolated using the NucleoSpin Plasmid EasyPure kit (Macherey-Nagel GmbH & Co.) and sent to Plasmidsaurus Inc. for long-read sequencing with Oxford Nanopore Technology.

### Statistical analysis

Data generated by the microplate readers were analyzed using the QurvE software tool^49^ to calculate maximum growth rates. All cultivations were performed in biological triplicates unless otherwise specified. All other data analysis was performed in R (v4.5.0, R Core Team) using RStudio (Posit Software, PBC). P-values were calculated using the limma package (v3.64.3) with moderated two-sided t-test.

## Acknowledgements

We acknowledge the DTU Proteomics Core for performing proteomics analysis. We thank Nicolas Thilo Wirth and Hanna Müller-Esparza for sharing strains and plasmids used in this study. This work was supported by the Novo Nordisk Foundation through grant NNF20CC0035580 and NNF24SA0100980.

## Authors contribution

**Mariana Arango Saavedra:** conceptualization, methodology, formal analysis, investigation, writing – original draft, writing – reviewing & editing, visualization; **Sara Grassi:** investigation, formal analysis; **Magnus G Jespersen**: formal analysis; **Catarina Rocha**: investigation, formal analysis; **Viji Kandasamy**: supervision, resources; **Lars Keld Nielsen**: conceptualization, supervision, funding acquisition; **Pablo Ivan Nikel**: conceptualization, supervision, resources; **Stefano Donati**: conceptualization, methodology, writing – original draft, writing – reviewing & editing, supervision, project administration.

## References

(1) Keasling, J. D. Manufacturing Molecules through Metabolic Engineering. Science 2010, 330 (6009), 1355–1358. 10.1126/science.1193990.

(2) Fisher, A. K.; Freedman, B. G.; Bevan, D. R.; Senger, R. S. A Review of Metabolic and Enzymatic Engineering Strategies for Designing and Optimizing Performance of Microbial Cell Factories. Computational and Structural Biotechnology Journal 2014, 11 (18), 91–99. 10.1016/j.csbj.2014.08.010.

(3) Chaudhary, R.; Nawaz, A.; Fouillaud, M.; Dufossé, L.; Haq, I. ul; Mukhtar, H.; Chaudhary, R.; Nawaz, A.; Fouillaud, M.; Dufossé, L.; Haq, I. ul; Mukhtar, H. Microbial Cell Factories: Biodiversity, Pathway Construction, Robustness, and Industrial Applicability. Microbiology Research 2024, 15 (1), 247–272. 10.3390/microbiolres15010018.

(4) Lee, S. Y.; Kim, H. U. Systems Strategies for Developing Industrial Microbial Strains. Nat Biotechnol 2015, 33 (10), 1061–1072. 10.1038/nbt.3365.

(5) Ko, Y.-S.; Woong Kim, J.; An Lee, J.; Han, T.; Bae Kim, G.; Eum Park, J.; Yup Lee, S. Tools and Strategies of Systems Metabolic Engineering for the Development of Microbial Cell Factories for Chemical Production. Chemical Society Reviews 2020, 49 (14), 4615–4636. 10.1039/D0CS00155D.

(6) Yeom, J.; Park, J. S.; Jung, S.-W.; Lee, S.; Kwon, H.; Yoo, S. M. High-Throughput Genetic Engineering Tools for Regulating Gene Expression in a Microbial Cell Factory. Critical Reviews in Biotechnology 2023, 43 (1), 82–99. 10.1080/07388551.2021.2007351.

(7) Carthew, R. W. Gene Regulation and Cellular Metabolism: An Essential Partnership. Trends Genet 2021, 37 (4), 389–400. 10.1016/j.tig.2020.09.018.

(8) Liu, Z.; Dong, H.; Cui, Y.; Cong, L.; Zhang, D. Application of Different Types of CRISPR/Cas-Based Systems in Bacteria. Microb Cell Fact 2020, 19 (1), 172. 10.1186/s12934-020-01431-z.

(9) Bikard, D.; Jiang, W.; Samai, P.; Hochschild, A.; Zhang, F.; Marraffini, L. A. Programmable Repression and Activation of Bacterial Gene Expression Using an Engineered CRISPR-Cas System. Nucleic Acids Res 2013, 41 (15), 7429–7437. 10.1093/nar/gkt520.

(10) Vigouroux, A.; Bikard, D. CRISPR Tools To Control Gene Expression in Bacteria. Microbiol Mol Biol Rev 2020, 84 (2), e00077–19. 10.1128/MMBR.00077-19.

(11) Frusteri Chiacchiera, A.; Casanova, M.; Bellato, M.; Piazza, A.; Migliavacca, R.; Batt, G.; Magni, P.; Pasotti, L. Harnessing CRISPR Interference to Resensitize Laboratory Strains and Clinical Isolates to Last Resort Antibiotics. Sci Rep 2025, 15 (1), 261. 10.1038/s41598-024-81989-5.

(12) Donati, S.; Kuntz, M.; Pahl, V.; Farke, N.; Beuter, D.; Glatter, T.; Gomes-Filho, J. V.; Randau, L.; Wang, C. Y.; Link, H. Multi-Omics Analysis of CRISPRi-Knockdowns Identifies Mechanisms That Buffer Decreases of Enzymes in E. Coli Metabolism. Cell Systems 2021, 12 (1), 56-67.e6. 10.1016/j.cels.2020.10.011.

(13) Fenster, J. A.; Werner, A. Z.; Tay, J. W.; Gillen, M.; Schirokauer, L.; Hill, N. C.; Watson, A.; Ramirez, K. J.; Johnson, C. W.; Beckham, G. T.; Cameron, J. C.; Eckert, C. A. Dynamic and Single Cell Characterization of a CRISPR-Interference Toolset in Pseudomonas Putida KT2440 for β-Ketoadipate Production from p-Coumarate. Metabolic Engineering Communications 2022, 15. 10.1016/j.mec.2022.e00204.

(14) Yao, L.; Shabestary, K.; Björk, S. M.; Asplund-Samuelsson, J.; Joensson, H. N.; Jahn, M.; Hudson, E. P. Pooled CRISPRi Screening of the Cyanobacterium Synechocystis Sp PCC 6803 for Enhanced Industrial Phenotypes. Nature Communications 2020, 11 (1). 10.1038/s41467-020-15491-7.

(15) Miao, R.; Jahn, M.; Shabestary, K.; Peltier, G.; Hudson, E. P. CRISPR Interference Screens Reveal Growth–Robustness Tradeoffs in Synechocystis Sp. PCC 6803 across Growth Conditions. The Plant Cell 2023. 10.1093/plcell/koad208.

(16) Carruthers, D. N.; Kinnunen, P. C.; Li, Y.; Chen, Y.; Gin, J. W.; Yunus, I. S.; Galliard, W. R.; Tan, S.; Radivojevic, T.; Adams, P. D.; Singh, A. K.; Sustarich, J.; Petzold, C. J.; Mukhopadhyay, A.; Garcia Martin, H.; Lee, T. S. Automation and Machine Learning Drive Rapid Optimization of Isoprenol Production in Pseudomonas Putida. Nat Commun 2025, 16 (1), 11489. 10.1038/s41467-025-66304-8.

(17) Yang, C.-C.; Deshpande, A. J.; Jackson, M.; Adams, P. D.; Pasquale, E. B.; Murad, R.; Yin, J.-A.; Wu, Y.; Beketova, A.; Huang, C.-T. Automation of High-Throughput Workflow for Arrayed CRISPR Activation Library Screening. bioRxiv 2025, 2025.11.10.687722. 10.1101/2025.11.10.687722.

(18) Steel, H.; Habgood, R.; Kelly, C.; Papachristodoulou, A. Chi.Bio: An Open-Source Automated Experimental Platform for Biological Science Research. 2019. 10.1101/796516.

(19) Pioreactor. Pioreactor Inc. https://pioreactor.com/en-dk (accessed 2025-10-07).

(20) Cui, L.; Vigouroux, A.; Rousset, F.; Varet, H.; Khanna, V.; Bikard, D. A CRISPRi Screen in E. Coli Reveals Sequence-Specific Toxicity of dCas9. Nat Commun 2018, 9 (1), 1912. 10.1038/s41467-018-04209-5.

(21) Rostain, W.; Grebert, T.; Vyhovskyi, D.; Pizarro, P. T.; Bellingen, G. T. V.; Cui, L.; Bikard, D. Cas9 Off-Target Binding to the Promoter of Bacterial Genes Leads to Silencing and Toxicity. Nucleic Acids Research 2023, 51 (7), 3485–3496. 10.1093/nar/gkad170.

(22) Wirth, N. T.; Kozaeva, E.; Nikel, P. I. Accelerated Genome Engineering of Pseudomonas Putida by I-SceI―mediated Recombination and CRISPR-Cas9 Counterselection. Microbial Biotechnology 2020, 13 (1), 233–249. 10.1111/1751-7915.13396.

(23) Scribani-Rossi, C.; Molina-Henares, M. A.; Espinosa-Urgel, M.; Rinaldo, S. Exploring the Metabolic Response of Pseudomonas Putida to L-Arginine; 2024. 10.1007/5584_2024_797.

(24) Barrientos-Moreno, L.; Molina-Henares, M. A.; Ramos-González, M. I.; Espinosa-Urgel, M. Arginine as an Environmental and Metabolic Cue for Cyclic Diguanylate Signalling and Biofilm Formation in Pseudomonas Putida. Sci Rep 2020, 10 (1), 13623. 10.1038/s41598-020-70675-x.

(25) Barrientos-Moreno, L.; Molina-Henares, M. A.; Ramos-González, M. I.; Espinosa-Urgel, M. Role of the Transcriptional Regulator ArgR in the Connection between Arginine Metabolism and C-Di-GMP Signaling in Pseudomonas Putida. Applied and Environmental Microbiology 2022, 88 (7). 10.1128/aem.00064-22.

(26) Nishijyo, T.; Park, S.-M.; Lu, C.-D.; Itoh, Y.; Abdelal, A. T. Molecular Characterization and Regulation of an Operon Encoding a System for Transport of Arginine and Ornithine and the ArgR Regulatory Protein in Pseudomonas Aeruginosa. Journal of Bacteriology 1998, 180 (21), 5559–5566. 10.1128/jb.180.21.5559-5566.1998.

(27) Molina-Henares, M. A.; Ramos-González, M. I.; Rinaldo, S.; Espinosa-Urgel, M. Gene Expression Reprogramming of Pseudomonas Alloputida in Response to Arginine through the Transcriptional Regulator ArgR. Microbiology (Reading) 2024, 170 (3), 001449. 10.1099/mic.0.001449.

(28) Johnson, C. W.; Abraham, P. E.; Linger, J. G.; Khanna, P.; Hettich, R. L.; Beckham, G. T. Eliminating a Global Regulator of Carbon Catabolite Repression Enhances the Conversion of Aromatic Lignin Monomers to Muconate in Pseudomonas Putida KT2440. Metabolic Engineering Communications 2017, 5, 19–25. 10.1016/j.meteno.2017.05.002.

(29) Shrestha, S.; Awasthi, D.; Chen, Y.; Gin, J.; Petzold, C. J.; Adams, P. D.; Simmons, B. A.; Singer, S. W. Simultaneous Carbon Catabolite Repression Governs Sugar and Aromatic Co-Utilization in Pseudomonas Putida M2. Applied and Environmental Microbiology 2023, 89 (10), e00852–23. 10.1128/aem.00852-23.

(30) Batianis, C.; Kozaeva, E.; Damalas, S. G.; Martín-Pascual, M.; Volke, D. C.; Nikel, P. I.; Martins dos Santos, V. A. P. An Expanded CRISPRi Toolbox for Tunable Control of Gene Expression in Pseudomonas Putida. Microbial Biotechnology 2020, 13 (2), 368–385. 10.1111/1751-7915.13533.

(31) Kim, S. K.; Yoon, P. K.; Kim, S.-J.; Woo, S.-G.; Rha, E.; Lee, H.; Yeom, S.-J.; Kim, H.; Lee, D.-H.; Lee, S.-G. CRISPR Interference-Mediated Gene Regulation in Pseudomonas Putida KT2440. Microbial Biotechnology 2020, 13 (1), 210–221. 10.1111/1751-7915.13382.

(32) Sun, J.; Wang, Q.; Jiang, Y.; Wen, Z.; Yang, L.; Wu, J.; Yang, S. Genome Editing and Transcriptional Repression in Pseudomonas Putida KT2440 via the Type II CRISPR System. Microb Cell Fact 2018, 17 (1), 41. 10.1186/s12934-018-0887-x.

(33) Czajka, J. J.; Banerjee, D.; Eng, T.; Menasalvas, J.; Yan, C.; Munoz, N. M.; Poirier, B. C.; Kim, Y.-M.; Baker, S. E.; Tang, Y. J.; Mukhopadhyay, A. Tuning a High Performing Multiplexed-CRISPRi Pseudomonas Putida Strain to Further Enhance Indigoidine Production. Metabolic Engineering Communications 2022, 15, e00206. 10.1016/j.mec.2022.e00206.

(34) Gauttam, R.; Mukhopadhyay, A.; Simmons, B. A.; Singer, S. W. Development of Dual-Inducible Duet-Expression Vectors for Tunable Gene Expression Control and CRISPR Interference-Based Gene Repression in Pseudomonas Putida KT2440. Microbial Biotechnology 2021, 14 (6), 2659–2678. 10.1111/1751-7915.13832.

(35) Jeske, M.; Altenbuchner, J. The Escherichia Coli Rhamnose Promoter rhaPBADis in Pseudomonas Putida KT2440 Independent of Crp–cAMP Activation. Appl Microbiol Biotechnol 2010, 85 (6), 1923–1933. 10.1007/s00253-009-2245-8.

(36) Meisner, J.; Goldberg, J. B. The Escherichia Coli rhaSR-PrhaBAD Inducible Promoter System Allows Tightly Controlled Gene Expression over a Wide Range in Pseudomonas Aeruginosa. Applied and Environmental Microbiology 2016, 82 (22), 6715–6727. 10.1128/AEM.02041-16.

(37) Sonnabend, M. S.; Klein, K.; Beier, S.; Angelov, A.; Kluj, R.; Mayer, C.; Groß, C.; Hofmeister, K.; Beuttner, A.; Willmann, M.; Peter, S.; Oberhettinger, P.; Schmidt, A.; Autenrieth, I. B.; Schütz, M.; Bohn, E. Identification of Drug Resistance Determinants in a Clinical Isolate of Pseudomonas Aeruginosa by High-Density Transposon Mutagenesis. Antimicrobial Agents and Chemotherapy 2020, 64 (3), 10.1128/aac.01771-19. https://doi.org/10.1128/aac.01771-19.

(38) Sydow, A.; Pannek, A.; Krieg, T.; Huth, I.; Guillouet, S. E.; Holtmann, D. Expanding the Genetic Tool Box for Cupriavidus Necator by a Stabilized L-Rhamnose Inducible Plasmid System. Journal of Biotechnology 2017, 263, 1–10. 10.1016/j.jbiotec.2017.10.002.

(39) Cardona, S. T.; Valvano, M. A. An Expression Vector Containing a Rhamnose-Inducible Promoter Provides Tightly Regulated Gene Expression in Burkholderia Cenocepacia. Plasmid 2005, 54 (3), 219–228. 10.1016/j.plasmid.2005.03.004.

(40) Calero, P.; Jensen, S. I.; Nielsen, A. T. Broad-Host-Range ProUSER Vectors Enable Fast Characterization of Inducible Promoters and Optimization of p-Coumaric Acid Production in Pseudomonas Putida KT2440. ACS Synth. Biol. 2016, 5 (7), 741–753. 10.1021/acssynbio.6b00081.

(41) Tang, B.; Sitomer, A.; Jackson, T. Population Dynamics and Competition in Chemostat Models with Adaptive Nutrient Uptake. J Math Biol 1997, 35 (4), 453–479. 10.1007/s002850050061.

(42) Piper, M. D. W.; Daran-Lapujade, P.; Bro, C.; Regenberg, B.; Knudsen, S.; Nielsen, J.; Pronk, J. T. Reproducibility of Oligonucleotide Microarray Transcriptome Analyses. An Interlaboratory Comparison Using Chemostat Cultures of Saccharomyces Cerevisiae. J Biol Chem 2002, 277 (40), 37001–37008. 10.1074/jbc.M204490200.

(43) Barrientos-Moreno, L.; Molina-Henares, M. A.; Pastor-García, M.; Ramos-González, M. I.; Espinosa-Urgel, M. Arginine Biosynthesis Modulates Pyoverdine Production and Release in Pseudomonas Putida as Part of the Mechanism of Adaptation to Oxidative Stress. Journal of Bacteriology 2019, 201 (22), 10.1128/jb.00454-19. https://doi.org/10.1128/jb.00454-19.

(44) Lambertsen, L.; Sternberg, C.; Molin, S. Mini-Tn7 Transposons for Site-Specific Tagging of Bacteria with Fluorescent Proteins. Environmental Microbiology 2004, 6 (7), 726–732. 10.1111/j.1462-2920.2004.00605.x.

(45) Cavaleiro, A. M.; Kim, S. H.; Seppälä, S.; Nielsen, M. T.; Nørholm, M. H. H. Accurate DNA Assembly and Genome Engineering with Optimized Uracil Excision Cloning. ACS Synthetic Biology 2015, 4 (9), 1042–1046. 10.1021/acssynbio.5b00113.

(46) Rocha, C.; Pinto, S. P.; Jensen, S. I.; Nielsen, L. K.; Donati, S. A Robust Isotope Ratio LC-MS/MS Workflow for High-Throughput Metabolic Profiling of Bacteria. bioRxiv August 12, 2025, p 2025.08.08.669109. 10.1101/2025.08.08.669109.

(47) Gurdo, N.; Taylor Parkins, S. K.; Fricano, M.; Wulff, T.; Nielsen, L. K.; Nikel, P. I. Protocol for Absolute Quantification of Proteins in Gram-Negative Bacteria Based on QconCAT-Based Labeled Peptides. STAR Protocols 2023, 4 (1), 102060. 10.1016/j.xpro.2023.102060.

(48) Taylor Parkins, S. K. Computational Tools and Analytical Methods for Effective Metabolic Engineering of Microbial Cell Factories; Technical University of Denmark: Kongens Lyngby, 2023.

(49) Wirth, N. T.; Funk, J.; Donati, S.; Nikel, P. I. QurvE: User-Friendly Software for the Analysis of Biological Growth and Fluorescence Data. Nat Protoc 2023, 18 (8), 2401–2403. 10.1038/s41596-023-00850-7.

