## Supplemental materials for "Automated mini-bioreactors reveal the temporal dynamics and multi-omics responses of CRISPRi knockdowns in *Pseudomonas putida*"

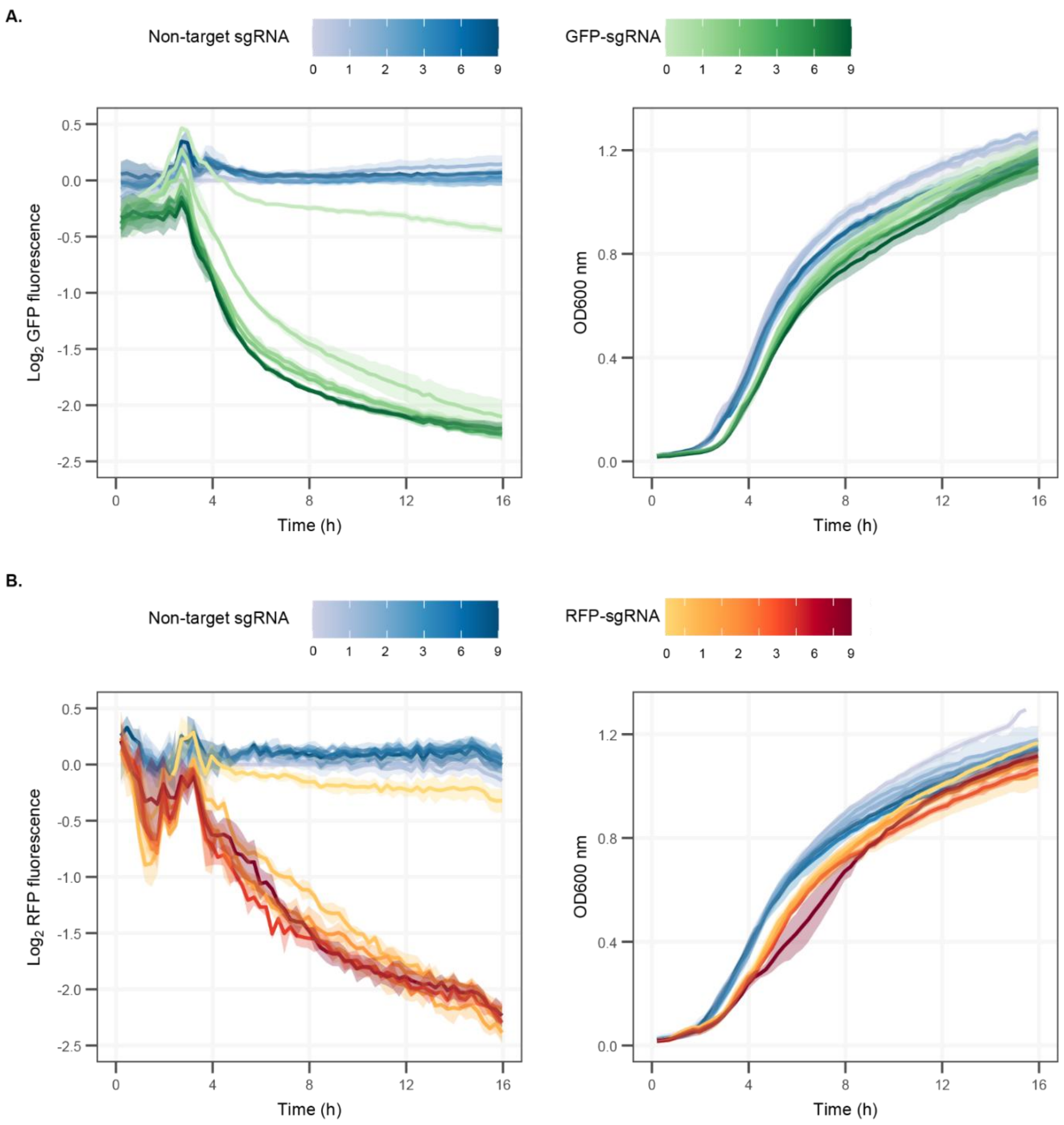


**Fig. S 1.** Effect of different concentrations of rhamnose on fluorescence repression and growth of *P. putida* KT2440 with genomically integrated GFP and RFP. **(A)** Fluorescence and growth profiles of GFP knockdown. **(B)**. Fluorescence and growth profiles of RFP knockdown. Fluorescence is normalized to OD and to fluorescent expression of an un-induced sg-RNA Control strain

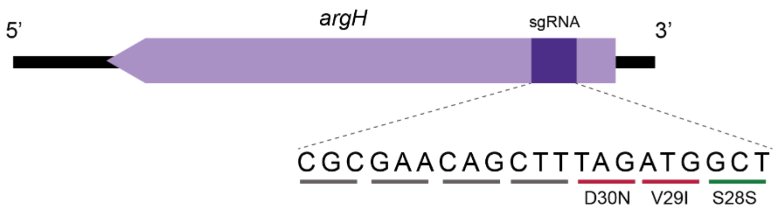


**Fig. S 2.** DNA sequence of argH and its target sgRNA. Point mutations are in bold, and synonymous and non-synonymous mutations are indicated in red and green, respectively.

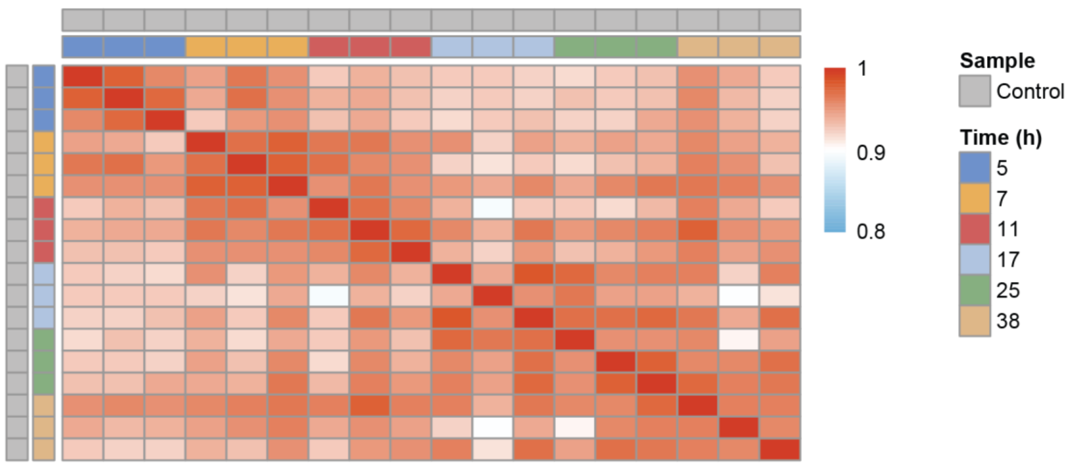


**Fig. S 3.** Heatmap showing pairwise Pearson correlation coefficients of proteomics profiles from cells harboring a sgRNA targeting *argH* under uninduced conditions (Control). Comparisons are made between individual replicates of the Control over time.

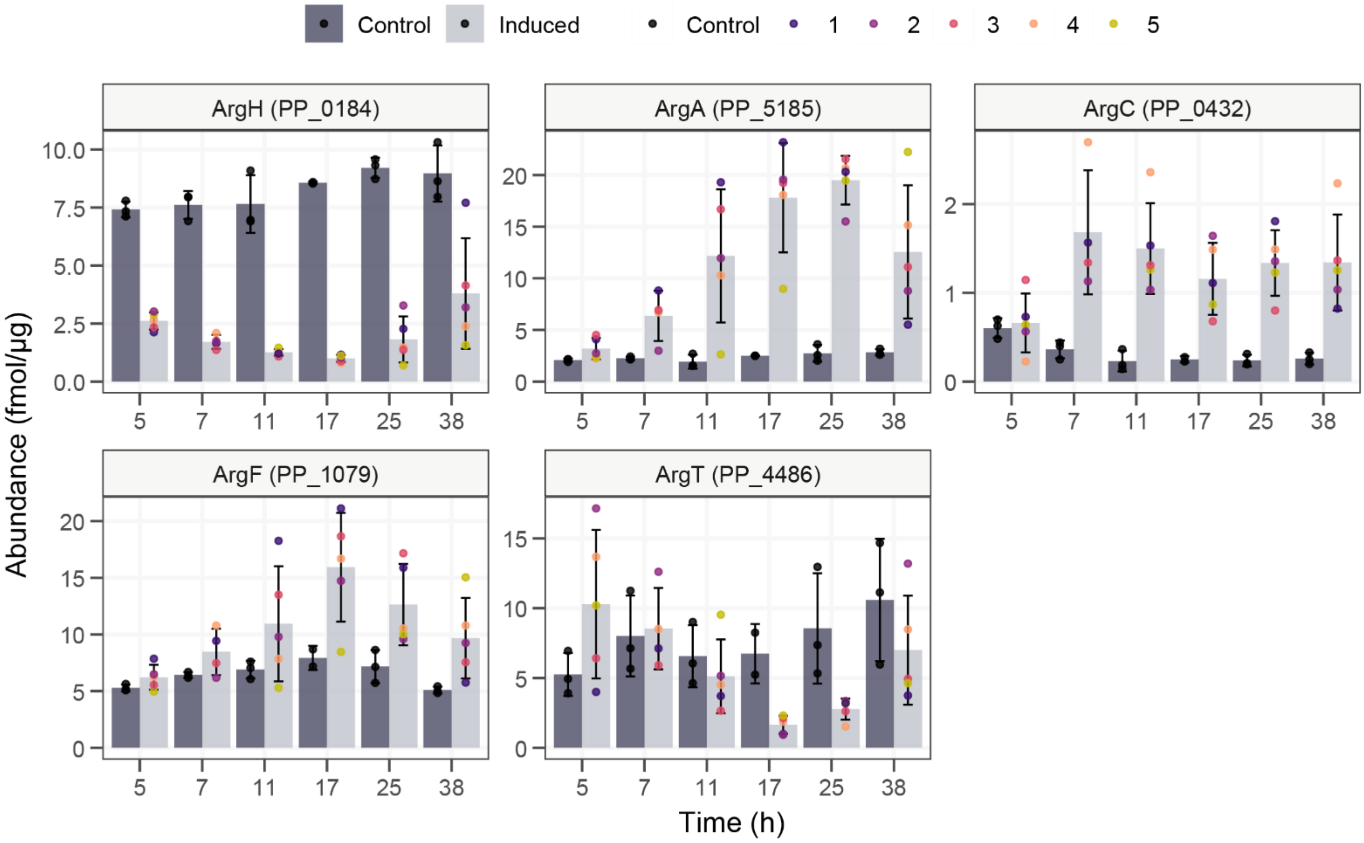


**Fig. S 4.** Concentrations of enzymes involved in arginine metabolism for *argH* knockdown over time. The uninduced strain carrying the sgRNA is referred to as Control. Individual replicates for the induced conditions are numbered according to their growth recovery capacity, from fastest to slowest. The bar plots represent mean values of three biological replicates for the control and five for the induced, with error bars indicating the standard deviation. Only enzymes that had a log2(foldchange) > 1 at 17 hours of induction are shown. Locus tag for each protein is indicated in parenthesis. Levels of statistical significance calculated with Wilcoxon t-test are shown as ****$p < 0.0001$, ***$p < 0.001$, **$p < 0.01$, *$p < 0.05$.

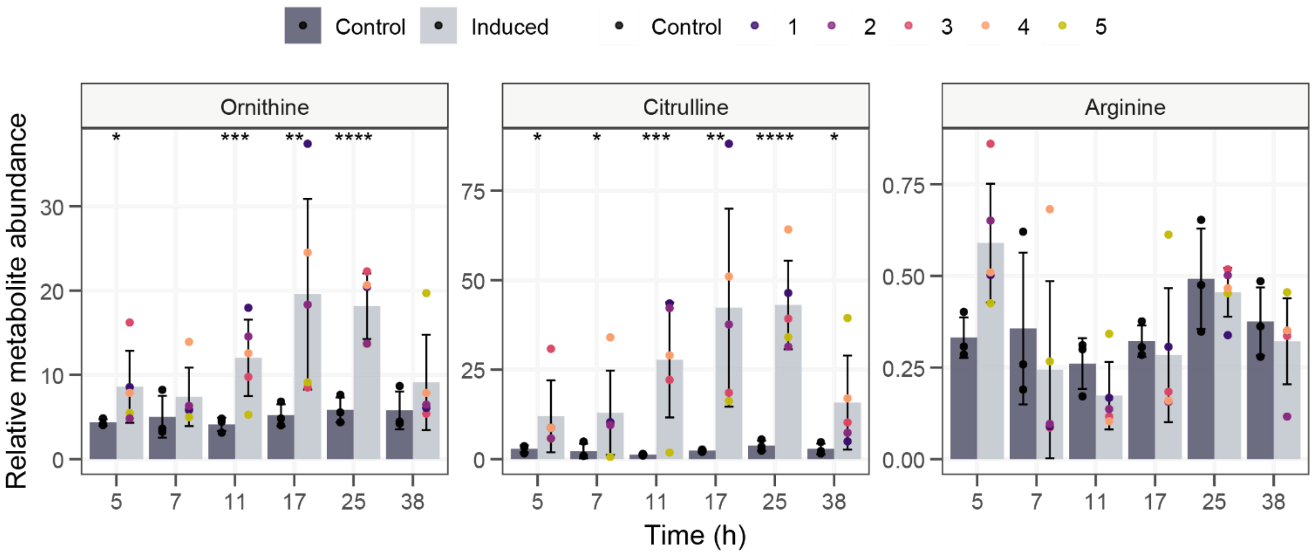


**Fig. S 5.** Relative abundances of intracellular ornithine, citrulline and arginine in an *argH* knockdown over time. The uninduced strain carrying the sgRNA is referred to as Control. Individual replicates for the induced conditions are numbered according to their growth recovery capacity, from fastest to slowest. The bar plots represent mean values of three biological replicates for the control and five for the induced, with error bars indicating the standard deviation. Levels of statistical significance calculated with Wilcoxon t-test are shown as **p < 0.01, *p < 0.05.

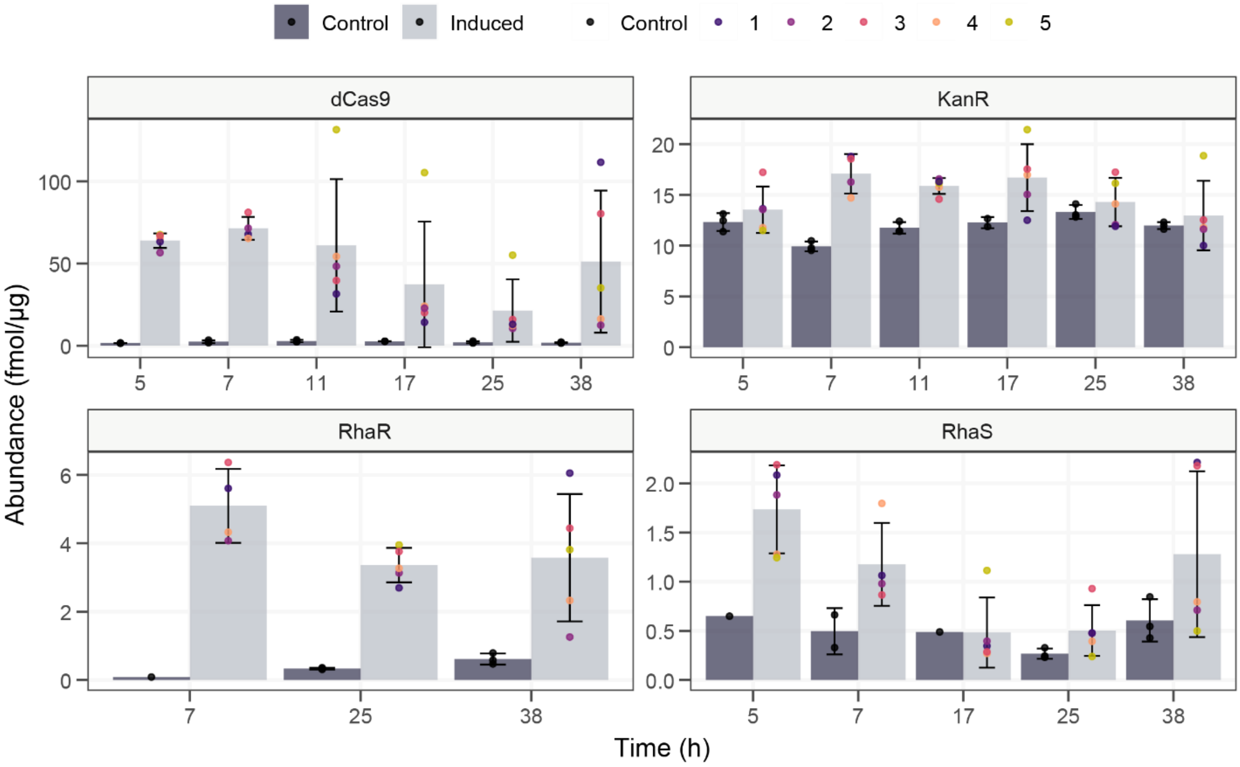


**Fig. S 6.** Concentration of proteins comprising the CRISPRi system in *argH* knockdown over time. The uninduced strain carrying the sgRNA is referred to as Control. Individual replicates for the induced conditions are numbered according to their growth recovery capacity, from fastest to slowest. The bar plots represent mean values of three biological replicates for the control and five for the induced, with error bars indicating the standard deviation. Levels of statistical significance calculated with Wilcoxon t-test are shown as ****p < 0.0001, ***p < 0.001, **p < 0.01, *p < 0.05. dCas9: deactivated Cas9; KanR: protein encoded by the kanamycin resistant gene; RhaR and RhaS: transcriptional regulators responsible for the activation of the rhamnose inducible promoter.

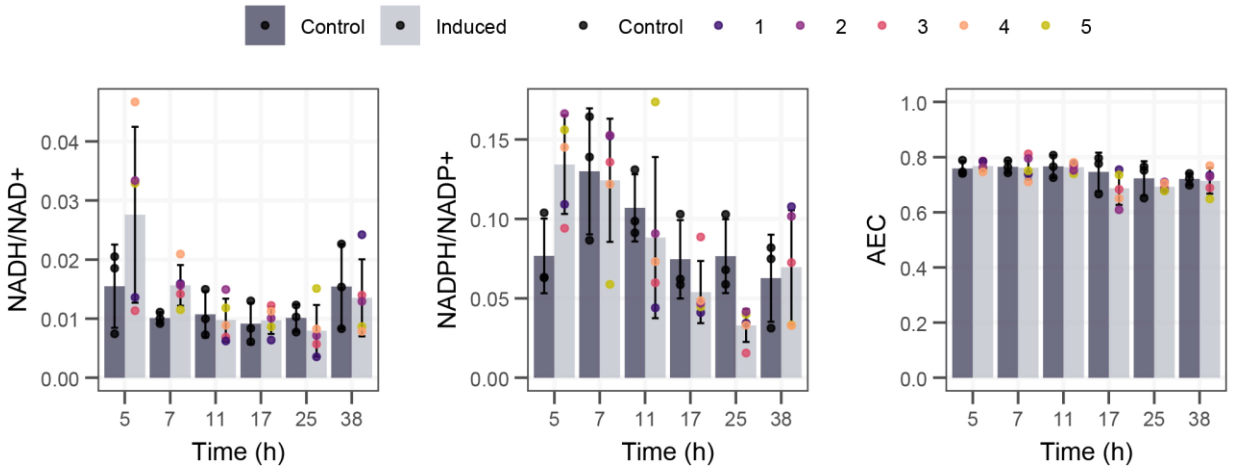


**Fig. S 7.** Oxidative balances and energy charge ratios for *argH* knockdown over time. The oxidative balances are showing relative abundances, while the Adenylate Energy Change (AEC) shows the mole fraction and was calculated as [atp]+0.5[adp] / [atp]+[adp]+[amp]. The uninduced strain carrying the sgRNA is referred to as Control. Individual replicates for the induced conditions are numbered according to their growth recovery capacity, from fastest to slowest. The bar plots represent mean values of three biological replicates for the control and five for the induced, with error bars indicating the standard deviation.

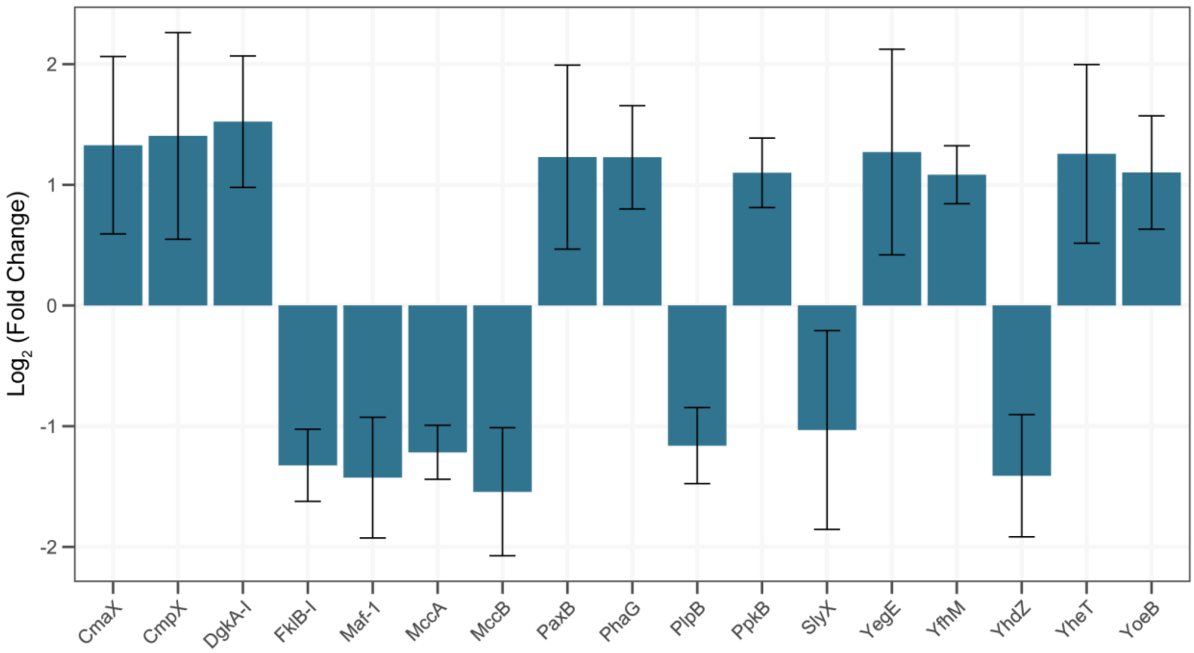


**Fig. S 8.** $Log_2$ fold change of a subset of significantly upregulated and downregulated proteins (p-value < 0.05) in *argH* knockdown after 17 hours of CRISPRi induction compared to non-induced control. Proteins shown belong to the 'Other' functional category.

**Table S 1**. Bacterial strains used in this study.

| Strain Name | Description | Source |
|---|---|---|
| NEB-beta10 | *E. coli* with genotype: *Δ(ara-leu) 7697 araD139 fhuA ΔlacX74 galK16 galE15 e14- ϕ80dlacZΔM15 recA1 relA1 endA1 nupG rpsL ($Str^R$) rph spoT1 Δ(mrr-hsdRMS-mcrBC).* | New England BioLabs Inc. |
| OneShot TOP10 | *E. coli* with genotype: F⁻*mcr*A Δ(*mrr-hsd*RMS-*mcr*BC) ф80*lac*ZΔM15 Δ*lac*X74 *rec*A1 *ara*D139 Δ(*ara-leu*)7697 *gal*U *gal*K λ–*rps*L($Str^R$) *end*A1 *nup*G. | Thermo Fisher Scientific Inc. |
| Dh5α | *E. coli* with genotype: *fhuA2::IS2 Δ(mmuP-mhpD)169 ΔphoA8 glnX44 ϕ80d[ΔlacZ58(M15)] rfbD1 gyrA96 luxS11 recA1 endA1 rphWT thiE1 hsdR17.* | New England BioLabs Inc. |
| KT2440 | *Pseudomonas putida* KT2440 strain. | Bagdasarian et al.[1] |
| KT·LPG | Derivative of KT2440 carrying $P_{14g}$(*BCD2*)→*msfGFP* element integrated between *PP_0013* (*gyrB*) and *PP_5421.* | Wirth et al.[2] |
| KT·LPR | Derivative of KT2440 carrying a $P_{14g}$(*BCD2*)→*mRFP1* element integrated between *PP_0013* (*gyrB*) and *PP_5421.* | Wirth et al.[2] |
| KT2440_MAS | Derivative of KT2440 integrated with dCas9 cassette. | This study |
| KT·LPG_MAS | Derivative of KT·LPG integrated with dCas9 cassette. | This study |
| KT·LPR_MAS | Derivative of KT·LPR integrated with dCas9 cassette. | This study |

**Table S 2.** Plasmids used in this study.

| Plasmid | Description | Source |
|---|---|---|
| pJM220 | Mobilizable base vector with MCS; $Gm^R$ on mini-Tn*7*T; *rhaSR-PrhaBAD* inducible promoter. | Meisner et al.[3] |
| pMCRi | *SpdCas9* gene; non-targeting sgRNA spacer under EM7promoter; gRNA scaffold; $Sm^R$/$Sp^R$. | Batianis et al.[4] |
| pJM220_R1 | pJM220 derivative bearing the *SpdCas9* gene. | This study |
| pRK600 | Contains mating formation genes for conjugation RP4/RK2. | Keen et al.[5] |
| pUX-BF13 | TnsABCD transposase subunits for Tn7 gene insertion; $Amp^R$; $Gm^R$. | Bao et al.[6] |
| pSEVA2311 | Standard cloning vector; *oriV* (pBBR1); $Kan^R$. | Benedetti et al.[7] |
| pC1001 | Plasmid for CRISPRi containing non-targeting sgRNA spacer under EM7 promoter; sgRNA scaffold; T1 and T0 terminators | This study |

Antibiotic markers and abbreviations: MCS, multiple cloning site; SpdCas9, dCas9 from *Streptococcus pyogenes*; Gm, gentamycin; Sm, streptomycin; Sp, spectinomycin; Amp, ampicillin; Kan, kanamycin.

**Table S 3.** Oligos used in this study.

| Name | Sequence (5' → 3') | Description |
|---|---|---|
| OL100 | ACTAATCUAGAGTCGACCTGCAGGCA | Amplification of pJM220 for USER cloning, forward. |
| OL101 | ATAGCTGUTTCCTCTGCAGAGCACTA | Amplification of pJM220 for USER cloning, reverse. |
| OL102 | ACAGCTAUGGATAAGAAATACTCAATAGGCT | Amplification of dCas9 from pMCRi for USER cloning, forward. |
| OL103 | AGATTAGUCACCTCCTAGCTGACTCA | Amplification of dCas9 from pMCRi for USER cloning, reverse. |
| MS02 | GAACAACGAACTGCTGGAGA | Confirm Tn7 integration of dCas9 cassette downstream of *glmS* gene, forward. |

| | | |
|---|---|---|
| MS01 | CACAGCATAACTGGACTGATTTC | Confirm Tn7 integration of dCas9 cassette downstream of *glmS* gene, reverse. |
| OL197 | TTTGAGTCAGCTAGGAGGTG | Confirm Tn7 integration of dCas9 cassette upstream of PP_5408 gene, forward. |
| MS03 | AGAACGGAGCGATTCATCAG | Confirm Tn7 integration of dCas9 cassette upstream of PP_5408 gene, reverse. |
| MS012 | AATTCGAGCUCGGTACCCGG | Amplification of pSEVA2311 backbone for USER cloning, forward. |
| MS013 | ATTAATUAAAGGCATCAAATAAAACGAA | Amplification of pSEVA2311 backbone for USER cloning, reverse. |
| MS014 | AATTAAUGTTGACAATTAATCATCGGCATAGTAT | Amplification of sgRNA cassette from pMCRi for USER cloning, forward. |
| MS015 | AGCTCGAATUGCACCGACTCGGTGCCACT | Amplification of sgRNA cassette from pMCRi for USER cloning, reverse. |
| MS016 | AGCGTTCACCGACAAACA | Confirm sgRNA introduction into pC1001, forward. |
| MS017 | ACCGAGCGTTCTGAACAAA | Confirm sgRNA introduction into pC1001, reverse. |
| Control_Fwd | GAGACCCGAGACTGGTCTCA | non-targeting sgRNA. |
| Control_Rv | TGAGACCAGTCTCGGGTCTC | non-targeting sgRNA. |
| GFP_Fwd | GCGCGAACCAGGATCGGAACAACAC | sgRNA targeting *msfGFP.* |
| GFP_Rv | AAACGTGTTGTTCCGATCCTGGTTC | sgRNA targeting *msfGFP.* |
| RFP_Fwd | GCGCGCGGGTGTTTAACGTAAG | sgRNA targeting *mRFP.* |
| RFP_Rv | AAACCTTACGTTAAACACCCG | sgRNA targeting *mRFP.* |
| argG_Fwd | GCGCGTGAGAATCACCGAAGTATCA | sgRNA targeting *argG.* |
| argG_Rv | AAACTGATACTTCGGTGATTCTCA | sgRNA targeting *argG.* |
| argH_Fwd | GCGCGGCGCTTGTCGAAATCTACCG | sgRNA targeting *argH.* |
| argH_Rv | AAACCGGTAGATTTCGACAAGCGCC | sgRNA targeting *argH.* |
| pyrF_Fwd | GCGCGCTCACGGGTAGGGAAATCCA | sgRNA targeting *pyrF.* |
| pyrF_Rv | AAACTGGATTTCCCTACCCGTGAGC | sgRNA targeting *pyrF.* |
| zwf_Fwd | GCGCGGCAAGGTTCGACACTGATTG | sgRNA targeting *zwfA.* |
| zwf_Rv | AAACCAATCAGTGTCGAACCTTGCC | sgRNA targeting *zwfA.* |

**Table S 4.** Point mutations emerging in individual replicates of CRISPRi dynamic induction targeting *argH* after 42 hours of continuous cultivation. Percentages indicate the abundance of each mutation in the specified replicate.

| Position | Mutation | Replicate 1 | Replicate 2 | Replicate 3 | Replicate 4 | Replicate 5 | Annotation |
|---|---|---|---|---|---|---|---|
| 236735 | C→T | 5.79% | 14.33% | 0.00% | 0.00% | 0.00% | D30N (GAT→AAT) |
| 236738 | C→T | 10.95% | 5.48% | 0.00% | 41.47% | 0.00% | V29I (GTA→ATA) |
| 236739 | C→T | 0.00% | 8.80% | 0.00% | 40.22% | 0.00% | S28S (TCG→TCA) |

**Table S 5**. Proteins with significant changes in protein abundance for *argH* induced knockdown after 17 hours of CRISPRi induction compared to uninduced condition. The significant threshold was set at a p-value < 0.05 and $|\log_2(\text{fold change})| > 1$.

| Uniprot ID | Gene name | p-value | $log_2$FC | Category |
|---|---|---|---|---|
| Q88CF9 | *cyaA* | 0.00085784 | 3.4581094 | Signal Transduction, Regulatory and Sensory Systems |
| Q88HB2 | *nfrB* | 0.02387396 | 2.2445792 | Membrane and Transport Proteins |
| Q88F13 | *puuE* | 0.00073870 | 2.1436728 | Amino Acid Metabolism |
| Q88N61 | *ampG* | 0.01126784 | 2.1171994 | Membrane and Transport Proteins |
| Q88IY8 | *mcpP* | 0.01454809 | 2.0862814 | Signal Transduction, Regulatory and Sensory Systems |
| Q88N83 | *ftsL* | 6.11924807e-05 | 2.0823667 | Cell Structure and Division |
| Q88N45 | *mcpG* | 0.00160559 | 2.0153384 | Signal Transduction, Regulatory and Sensory Systems |
| Q88KR2 | *mrpD* | 0.00021397 | 2.0052695 | Membrane and Transport Proteins |
| Q88QK8 | *bfr-I* | 1.04203300e-06 | 1.8944359 | Stress Response, Redox and Detoxification |
| Q88HH2 | *vgrG-II* | 0.00676610 | 1.8842098 | Membrane and Transport Proteins |
| Q88NC0 | *moaC* | 0.00029209 | 1.8616594 | Cofactor, Vitamin and Nucleotide Metabolism |
| Q88QF4 | *dedA* | 0.01745546 | 1.7901636 | Membrane and Transport Proteins |
| Q88HA4 | *mexB* | 0.00746372 | 1.7524496 | Membrane and Transport Proteins |
| Q88RL6 | *metI* | 0.03874902 | 1.7439021 | Membrane and Transport Proteins |
| Q88L24 | *acnA-I* | 0.00041917 | 1.7091482 | Carbohydrate and Central Carbon Metabolism |
| Q88E10 | *mcpS* | 0.00556598 | 1.7049545 | Signal Transduction, Regulatory and Sensory Systems |
| Q88KP1 | *mcpA* | 0.00350772 | 1.7028964 | Signal Transduction, Regulatory and Sensory Systems |
| Q88HI5 | *cpxR* | 0.00050420 | 1.6832870 | Signal Transduction, Regulatory and Sensory Systems |
| Q88M99 | *cobD* | 0.00083588 | 1.6808522 | Cofactor, Vitamin and Nucleotide Metabolism |
| Q88NX1 | *bfr-II* | 4.21937349e-05 | 1.6379567 | Stress Response, Redox and Detoxification |
| Q88F06 | *yedI* | 0.01167778 | 1.6222619 | Stress Response, Redox and Detoxification |

| | | | | |
|---|---|---|---|---|
| Q88GG0 | *opdN* | 0.00184031 | 1.6156566 | Membrane and Transport Proteins |
| Q88RE6 | *algR* | 0.00059424 | 1.5950146 | Signal Transduction, Regulatory and Sensory Systems |
| Q88DC2 | *motA* | 0.00482234 | 1.5947024 | Membrane and Transport Proteins |
| Q88GZ6 | *quiA* | 0.01347259 | 1.5924866 | Aromatic Compound Metabolism |
| Q88M25 | *asnB* | 0.00111508 | 1.5738470 | Amino Acid Metabolism |
| Q88PC3 | *aroP-I* | 0.00967953 | 1.5706677 | Membrane and Transport Proteins |
| Q88RH5 | *scpC* | 0.00027818 | 1.5578946 | Cell Structure and Division |
| Q88NC7 | *algK* | 0.03329503 | 1.5563007 | Cell Structure and Division |
| Q88IP1 | *grx* | 0.00847539 | 1.5464373 | Stress Response, Redox and Detoxification |
| Q88LN6 | *fadE* | 0.02788484 | 1.5435645 | Stress Response, Redox and Detoxification |
| Q88MD7 | *dgkA-I* | 0.00460814 | 1.5229737 | Other |
| Q88E89 | *pxpA2* | 2.21053342e-05 | 1.5204248 | Amino Acid Metabolism |
| Q88C45 | *aspA* | 0.00080465 | 1.5157604 | Amino Acid Metabolism |
| Q88E23 | *mscL* | 0.00367476 | 1.5115134 | Membrane and Transport Proteins |
| Q88G79 | *ramA* | 0.02429320 | 1.4619116 | Signal Transduction, Regulatory and Sensory Systems |
| Q88RL0 | *fur* | 0.01494742 | 1.4427366 | Signal Transduction, Regulatory and Sensory Systems |
| Q88KY5 | *lolC* | 0.01615947 | 1.4409767 | Membrane and Transport Proteins |
| Q88MC3 | *gacS* | 0.02661793 | 1.4368441 | Signal Transduction, Regulatory and Sensory Systems |
| Q88L48 | *cmpX* | 0.02769826 | 1.4048797 | Other |
| P0A155 | *ahpF* | 0.00271895 | 1.4032515 | Stress Response, Redox and Detoxification |
| Q88KB4 | *cti* | 3.95177670e-05 | 1.3829288 | Stress Response, Redox and Detoxification |
| Q88NN4 | *galP-I* | 0.01046583 | 1.3769504 | Membrane and Transport Proteins |
| Q88CP1 | *cadA-III* | 0.00275618 | 1.3637392 | Amino Acid Metabolism |
| Q88PX4 | *lolB* | 0.00793121 | 1.3589939 | Cell Structure and Division |
| Q88EU4 | *fliL* | 0.00641245 | 1.3348890 | Membrane and Transport Proteins |
| Q88DG8 | *yhjG* | 0.01211931 | 1.3286804 | Membrane and Transport Proteins |

| | | | | |
|---|---|---|---|---|
| Q88L50 | *cmaX* | 0.00855508 | 1.3277121 | Other |
| Q88F54 | *pvdY* | 1.17897973e-05 | 1.3252830 | Cofactor, Vitamin and Nucleotide Metabolism |
| Q88F24 | *zipA* | 0.00052343 | 1.3194936 | Cell Structure and Division |
| Q88GE6 | *proC* | 0.00070746 | 1.3058547 | Amino Acid Metabolism |
| Q88KH3 | *ybbA* | 0.01207518 | 1.2984159 | Cofactor, Vitamin and Nucleotide Metabolism |
| Q88N93 | *petC* | 0.04274540 | 1.2886379 | Stress Response, Redox and Detoxification |
| Q88N84 | *rsmH* | 0.00050311 | 1.2768381 | Cofactor, Vitamin and Nucleotide Metabolism |
| Q88IS4 | *mqo3* | 0.00019290 | 1.2759023 | Carbohydrate and Central Carbon Metabolism |
| Q88IQ9 | *yefM* | 0.01071078 | 1.2732894 | Signal Transduction, Regulatory and Sensory Systems |
| Q88R99 | *oprE* | 0.00072201 | 1.2716136 | Membrane and Transport Proteins |
| Q88M14 | *yegE* | 0.02347091 | 1.2709841 | Other |
| Q88R13 | *ltaE* | 0.00088930 | 1.2571177 | Amino Acid Metabolism |
| Q88CR3 | *yheT* | 0.03102835 | 1.2565188 | Other |
| Q88E67 | *moaB-II* | 0.00014002 | 1.2553178 | Cofactor, Vitamin and Nucleotide Metabolism |
| Q88FS2 | *icd* | 0.00207248 | 1.2385037 | Carbohydrate and Central Carbon Metabolism |
| Q88EA1 | *aceK* | 0.00015602 | 1.2318172 | Carbohydrate and Central Carbon Metabolism |
| Q88N14 | *dctB* | 0.00492888 | 1.2299382 | Signal Transduction, Regulatory and Sensory Systems |
| Q88RG3 | *paxB* | 0.01962709 | 1.2291579 | Other |
| O85207 | *phaG* | 0.02369382 | 1.2276817 | Other |
| Q88RJ7 | *kinB* | 0.00084945 | 1.2081475 | Signal Transduction, Regulatory and Sensory Systems |
| Q88LW9 | *pgi1* | 0.01496737 | 1.1984255 | Carbohydrate and Central Carbon Metabolism |
| Q88D16 | *ubiJ* | 0.00049356 | 1.1914210 | Cofactor, Vitamin and Nucleotide Metabolism |
| Q88N85 | *mraZ* | 0.00266681 | 1.1900483 | Signal Transduction, Regulatory and Sensory Systems |

| | | | | |
|---|---|---|---|---|
| Q88PW1 | *pagL-I* | 0.00030863 | 1.1894578 | Cell Structure and Division |
| Q88F12 | *pucL* | 0.02157401 | 1.1730467 | Carbohydrate and Central Carbon Metabolism |
| Q88DL8 | *mrdA-II* | 0.00135041 | 1.1647400 | Cell Structure and Division |
| Q88E71 | *ybaO* | 0.00385677 | 1.1566919 | Signal Transduction, Regulatory and Sensory Systems |
| Q88FR2 | *cpo* | 0.00100387 | 1.1547896 | Stress Response, Redox and Detoxification |
| Q88DN0 | *lptE* | 0.00076594 | 1.1510423 | Membrane and Transport Proteins |
| Q88FN8 | *treZ* | 0.00302943 | 1.1496201 | Carbohydrate and Central Carbon Metabolism |
| Q88Q23 | *gdhA* | 0.00049510 | 1.1496113 | Amino Acid Metabolism |
| Q88RR7 | *tag* | 0.00329136 | 1.1457760 | Carbohydrate and Central Carbon Metabolism |
| A0A140FW35 | *rluC* | 0.00087673 | 1.1429002 | Cofactor, Vitamin and Nucleotide Metabolism |
| Q88N80 | *murF* | 6.65125967e-05 | 1.1408523 | Cell Structure and Division |
| Q88CS3 | *mtgA* | 0.00389658 | 1.1403784 | Cell Structure and Division |
| Q88N29 | *ttgR* | 0.01328328 | 1.1346263 | Signal Transduction, Regulatory and Sensory Systems |
| Q88P74 | *holC* | 0.04289789 | 1.1327610 | Cofactor, Vitamin and Nucleotide Metabolism |
| Q88PR3 | *cysZ* | 0.01892348 | 1.1312921 | Membrane and Transport Proteins |
| Q88D04 | *opgH* | 0.01360914 | 1.1261076 | Cell Structure and Division |
| Q88DD5 | *hflK* | 0.00088662 | 1.1041654 | Stress Response, Redox and Detoxification |
| Q88MY5 | *rnc* | 0.00044564 | 1.1035667 | Cofactor, Vitamin and Nucleotide Metabolism |
| Q88IR0 | *yoeB* | 0.00706331 | 1.1019964 | Other |
| Q88PY6 | *ppkB* | 0.01341970 | 1.0992992 | Other |
| Q88EV1 | *flhB* | 0.01155089 | 1.0897757 | Membrane and Transport Proteins |
| Q88GH9 | *glcC* | 0.02765348 | 1.0850486 | Signal Transduction, Regulatory and Sensory Systems |
| Q88QC4 | *yfhM* | 0.00058456 | 1.0831618 | Other |
| Q88DP9 | *fieF* | 0.02170455 | 1.0771586 | Membrane and Transport Proteins |

| | | | | |
|---|---|---|---|---|
| Q88IN0 | *csoR* | 0.00066178 | 1.0562127 | Signal Transduction, Regulatory and Sensory Systems |
| Q88I72 | *galE* | 0.00262072 | 1.0488045 | Carbohydrate and Central Carbon Metabolism |
| Q88FS8 | *ftsK* | 0.03857149 | 1.0446008 | Cell Structure and Division |
| Q88N92 | *sspA* | 0.00019144 | 1.0414987 | Stress Response, Redox and Detoxification |
| Q88N82 | *ftsI* | 0.00144510 | 1.0397661 | Cell Structure and Division |
| Q88R66 | *oprQ* | 0.00139270 | 1.0375692 | Membrane and Transport Proteins |
| Q88FN1 | *glgB* | 0.00022971 | 1.0372918 | Carbohydrate and Central Carbon Metabolism |
| Q88F14 | *pucM* | 0.00120548 | 1.0369541 | Cofactor, Vitamin and Nucleotide Metabolism |
| Q88MS5 | *cheB3* | 0.00106413 | 1.0348402 | Signal Transduction, Regulatory and Sensory Systems |
| Q88DC7 | *queG* | 0.00177027 | 1.0214749 | Cofactor, Vitamin and Nucleotide Metabolism |
| Q88JX7 | *galR* | 0.03173915 | 1.0158541 | Signal Transduction, Regulatory and Sensory Systems |
| Q88MR4 | *ppc* | 0.00060097 | 1.0111520 | Carbohydrate and Central Carbon Metabolism |
| Q88MY2 | *pdxJ* | 0.00053690 | 1.0076493 | Cofactor, Vitamin and Nucleotide Metabolism |
| Q88D12 | *tatB* | 0.02902146 | 1.0072820 | Membrane and Transport Proteins |
| Q88F73 | *qseB* | 0.00092243 | 1.0052340 | Signal Transduction, Regulatory and Sensory Systems |
| Q88FI4 | *fusB* | 0.00495316 | 1.0026525 | Stress Response, Redox and Detoxification |
| Q88L45 | *cobA* | 0.00240489 | -1.0005004 | Cofactor, Vitamin and Nucleotide Metabolism |
| Q88D27 | *hslU* | 0.00044546 | -1.0149835 | Stress Response, Redox and Detoxification |
| Q88MH9 | *tsf* | 0.00031846 | -1.0190076 | Cofactor, Vitamin and Nucleotide Metabolism |
| Q88I54 | *potF-II* | 0.00892877 | -1.0296218 | Membrane and Transport Proteins |
| Q88NJ9 | *slyX* | 0.04488822 | -1.0315132 | Other |
| Q88F00 | *ttuD* | 0.00666498 | -1.0496330 | Carbohydrate and Central Carbon Metabolism |

| | | | | |
|---|---|---|---|---|
| Q88MH7 | *frr* | 0.00688078 | -1.0535542 | Cofactor, Vitamin and Nucleotide Metabolism |
| Q88LE5 | *leuB* | 0.00015416 | -1.0576363 | Amino Acid Metabolism |
| Q88DU1 | *grpE* | 0.00051729 | -1.0596160 | Stress Response, Redox and Detoxification |
| Q88GX6 | *dpkA* | 0.00087963 | -1.0623949 | Amino Acid Metabolism |
| Q88IY3 | *azoR1* | 0.04268278 | -1.0907949 | Stress Response, Redox and Detoxification |
| Q88FM3 | *liuC* | 0.00730603 | -1.0914264 | Carbohydrate and Central Carbon Metabolism |
| Q88R97 | *ssuE* | 0.00049109 | -1.1156676 | Stress Response, Redox and Detoxification |
| Q88IS0 | *nspC* | 0.00025250 | -1.1175739 | Amino Acid Metabolism |
| Q88DG2 | *livF-II* | 0.00077370 | -1.1178122 | Membrane and Transport Proteins |
| Q88LE2 | *asd* | 0.00020101 | -1.1302502 | Amino Acid Metabolism |
| Q88CK9 | *sbp-II* | 0.00676111 | -1.1321541 | Membrane and Transport Proteins |
| Q88I38 | *benC* | 0.03134542 | -1.1355675 | Aromatic Compound Metabolism |
| Q88CZ9 | *hutI* | 0.03225966 | -1.1459291 | Amino Acid Metabolism |
| Q88P64 | *gcvH1* | 0.01520468 | -1.1467642 | Amino Acid Metabolism |
| Q88I37 | *benD* | 0.02091678 | -1.1541683 | Aromatic Compound Metabolism |
| Q88CL5 | *plpB* | 0.00075367 | -1.1612567 | Other |
| Q88C80 | *ridA* | 0.00190129 | -1.1661727 | Amino Acid Metabolism |
| Q88R42 | *hisA* | 0.00046887 | -1.1718520 | Amino Acid Metabolism |
| Q88HS3 | *paaJ* | 0.00558414 | -1.1770195 | Aromatic Compound Metabolism |
| Q88HL1 | *nikD* | 0.00064219 | -1.1840023 | Stress Response, Redox and Detoxification |
| Q88E69 | *moaA* | 0.00612834 | -1.1959031 | Cofactor, Vitamin and Nucleotide Metabolism |
| Q88P41 | *gltR-II* | 0.00351166 | -1.2013916 | Signal Transduction, Regulatory and Sensory Systems |
| Q88QN0 | *rplV* | 0.00222847 | -1.2021347 | Cell Structure and Division |
| Q88HA1 | *peaD* | 0.00076494 | -1.2115652 | Amino Acid Metabolism |
| Q88FM2 | *mccA* | 0.00041018 | -1.2159111 | Other |
| P0A122 | *fdxA* | 0.00095053 | -1.2192813 | Stress Response, Redox and Detoxification |

| | | | | |
|---|---|---|---|---|
| Q88GW6 | *garD* | 0.00302806 | -1.2212642 | Amino Acid Metabolism |
| Q88P37 | *gtsB* | 0.01076326 | -1.2629700 | Membrane and Transport Proteins |
| Q88MV6 | *rpsP* | 0.00144806 | -1.2705631 | Cell Structure and Division |
| Q88E47 | *hmgA* | 0.00018916 | -1.2968276 | Aromatic Compound Metabolism |
| Q88CZ6 | *hutU* | 0.00200855 | -1.3112982 | Amino Acid Metabolism |
| Q88N39 | *pcaF-I* | 0.00320867 | -1.3158886 | Aromatic Compound Metabolism |
| Q88Q14 | *fklB-I* | 0.00034696 | -1.3237120 | Other |
| Q88GX1 | *amaD* | 0.00228356 | -1.3263384 | Amino Acid Metabolism |
| Q88FS4 | *clpS* | 0.01175105 | -1.3340931 | Stress Response, Redox and Detoxification |
| Q88CN8 | *lgt* | 0.03962238 | -1.3549221 | Cell Structure and Division |
| Q88GJ9 | *alr* | 2.76576655e-05 | -1.3781695 | Amino Acid Metabolism |
| Q88PI2 | *yehX* | 0.00617427 | -1.3831141 | Membrane and Transport Proteins |
| Q88K16 | *iorA-I* | 0.00187343 | -1.4025680 | Stress Response, Redox and Detoxification |
| Q88NB2 | *yhdZ* | 0.00693096 | -1.4095439 | Other |
| Q88PB4 | *maf-1* | 0.00080149 | -1.4254688 | Other |
| Q88N56 | *groES* | 0.00023020 | -1.4418534 | Stress Response, Redox and Detoxification |
| Q88P44 | *gapA* | 5.85706252e-05 | -1.4556483 | Carbohydrate and Central Carbon Metabolism |
| Q88I39 | *benB* | 0.00289308 | -1.4639463 | Aromatic Compound Metabolism |
| Q88LE8 | *leuC* | 6.08537456e-05 | -1.5400849 | Amino Acid Metabolism |
| Q88FM4 | *mccB* | 0.00078558 | -1.5431581 | Other |
| Q88IU8 | *pvdQ* | 0.03091752 | -1.5479697 | Cofactor, Vitamin and Nucleotide Metabolism |
| Q88ND4 | *algF* | 0.00831437 | -1.5578282 | Cell Structure and Division |
| Q88M34 | *betX* | 0.01950949 | -1.5672076 | Membrane and Transport Proteins |
| Q88I35 | *catA-II* | 0.00117125 | -1.5675996 | Aromatic Compound Metabolism |
| Q88Q39 | *metB* | 0.00439151 | -1.5722676 | Amino Acid Metabolism |
| Q88LE7 | *leuD* | 0.00027139 | -1.5992215 | Amino Acid Metabolism |
| Q88PG8 | *dppA-I* | 0.00024844 | -1.6009943 | Membrane and Transport Proteins |
| Q88R36 | *gbdR* | 0.04134921 | -1.6450111 | Signal Transduction, Regulatory and Sensory Systems |

| | | | | |
|---|---|---|---|---|
| Q88K38 | *rbsB* | 0.00012204 | -1.6473411 | Membrane and Transport Proteins |
| Q88IB9 | *aroF-II* | 0.00235391 | -1.6481426 | Aromatic Compound Metabolism |
| Q88EQ1 | *bkdAB* | 0.01336871 | -1.6586364 | Amino Acid Metabolism |
| Q88JH0 | *pedH* | 0.04238586 | -1.6753610 | Stress Response, Redox and Detoxification |
| Q88GH7 | *glcE* | 0.00685412 | -1.6971675 | Carbohydrate and Central Carbon Metabolism |
| Q88E09 | *ggt* | 0.00026328 | -1.6996296 | Amino Acid Metabolism |
| Q88JV4 | *gabP-II* | 0.00075379 | -1.7401411 | Membrane and Transport Proteins |
| Q88FF2 | *mltD* | 0.01931609 | -1.7677020 | Cell Structure and Division |
| Q88MP8 | *cspA-I* | 4.80466941e-06 | -1.8615145 | Stress Response, Redox and Detoxification |
| Q88FX5 | *pcaI* | 0.00170145 | -1.8908043 | Aromatic Compound Metabolism |
| Q88E12 | *pcaH* | 0.00086764 | -1.9094024 | Aromatic Compound Metabolism |
| Q88PG6 | *dppA-II* | 0.00092373 | -1.9279569 | Membrane and Transport Proteins |
| Q88P35 | *gtsD* | 7.97548667e-06 | -1.9332874 | Membrane and Transport Proteins |
| Q88FM5 | *ivd* | 3.99614630e-05 | -1.9616696 | Amino Acid Metabolism |
| Q88HE4 | *gnuK* | 5.14160957e-06 | -1.9668732 | Carbohydrate and Central Carbon Metabolism |
| Q88GE1 | *syrB* | 1.65064767e-05 | -1.9798683 | Signal Transduction, Regulatory and Sensory Systems |
| Q88PG5 | *dppA-III* | 4.69371996e-05 | -2.0092429 | Membrane and Transport Proteins |
| Q88P38 | *gtsA* | 0.00017359 | -2.0186014 | Membrane and Transport Proteins |
| Q88P34 | *oprB-I* | 1.20659604e-05 | -2.0506522 | Membrane and Transport Proteins |
| Q88I40 | *benA* | 7.20984937e-05 | -2.0802311 | Aromatic Compound Metabolism |
| Q88GD8 | *aspC* | 9.68906504e-05 | -2.1294443 | Amino Acid Metabolism |
| Q88PG7 | *opdP* | 2.52379515e-05 | -2.1377971 | Membrane and Transport Proteins |
| Q88NR4 | *livK* | 0.00014807 | -2.1618484 | Membrane and Transport Proteins |
| Q88F47 | *ccoQ-I* | 0.02411741 | -2.1736269 | Stress Response, Redox and Detoxification |
| Q88GK6 | *catB* | 0.00138220 | -2.2137863 | Aromatic Compound Metabolism |
| Q88GJ2 | *aruR* | 2.69404368e-05 | -2.5219900 | Signal Transduction, Regulatory and Sensory Systems |
| Q88N33 | *galP-II* | 0.00075936 | -2.5411145 | Membrane and Transport Proteins |

| | | | | |
|---|---|---|---|---|
| Q88GK8 | *catA-I* | 0.00044376 | -2.5423007 | Aromatic Compound Metabolism |
| Q88NB5 | *yhdW* | 2.41341330e-07 | -2.7286308 | Membrane and Transport Proteins |

### Supplementary references


(1) Bagdasarian, M.; Lurz, R.; Rückert, B.; Franklin, F. C. H.; Bagdasarian, M. M.; Frey, J.; Timmis, K. N. Specific-Purpose Plasmid Cloning Vectors II. Broad Host Range, High Copy Number, RSF 1010-Derived Vectors, and a Host-Vector System for Gene Cloning in *Pseudomonas*. *Gene* **1981**, *16* (1), 237–247. https://doi.org/10.1016/0378-1119(81)90080-9.

(2) Wirth, N. T.; Kozaeva, E.; Nikel, P. I. Accelerated Genome Engineering of Pseudomonas Putida by I-SceI—mediated Recombination and CRISPR-Cas9 Counterselection. *Microbial Biotechnology* **2020**, *13* (1), 233–249. https://doi.org/10.1111/1751-7915.13396.

(3) Meisner, J.; Goldberg, J. B. The Escherichia Coli rhaSR-PrhaBAD Inducible Promoter System Allows Tightly Controlled Gene Expression over a Wide Range in Pseudomonas Aeruginosa. *Applied and Environmental Microbiology* **2016**, *82* (22), 6715–6727. https://doi.org/10.1128/AEM.02041-16.

(4) Batianis, C.; Kozaeva, E.; Damalas, S. G.; Martín-Pascual, M.; Volke, D. C.; Nikel, P. I.; Martins dos Santos, V. A. P. An Expanded CRISPRi Toolbox for Tunable Control of Gene Expression in Pseudomonas Putida. *Microbial Biotechnology* **2020**, *13* (2), 368–385. https://doi.org/10.1111/1751-7915.13533.

(5) Keen, N. T.; Tamaki, S.; Kobayashi, D.; Trollinger, D. Improved Broad-Host-Range Plasmids for DNA Cloning in Gram-Negative Bacteria. *Gene* **1988**, *70* (1), 191–197. https://doi.org/10.1016/0378-1119(88)90117-5.

(6) Bao, Y.; Lies, D. P.; Fu, H.; Roberts, G. P. An Improved Tn7-Based System for the Single-Copy Insertion of Cloned Genes into Chromosomes of Gram-Negative Bacteria. *Gene* **1991**, *109* (1), 167–168. https://doi.org/10.1016/0378-1119(91)90604-A.

(7) Benedetti, I.; Nikel, P. I.; de Lorenzo, V. Data on the Standardization of a Cyclohexanone-Responsive Expression System for Gram-Negative Bacteria. *Data in Brief* **2016**, *6*, 738–744. https://doi.org/10.1016/j.dib.2016.01.022.
